# Computational remodeling of an enzyme conformational landscape for altered substrate selectivity

**DOI:** 10.1101/2022.09.16.508321

**Authors:** Antony D. St-Jacques, Joshua M. Rodriguez, Matthew G. Eason, Scott M. Foster, Safwat T. Khan, Adam M. Damry, Natalie K. Goto, Michael C. Thompson, Roberto A. Chica

## Abstract

Structural plasticity of enzymes dictates their function. Yet, our ability to rationally remodel enzyme conformational landscapes to tailor catalytic properties remains limited. Here, we report a computational procedure for tuning conformational landscapes that is based on multistate design. Using this method, we redesigned the conformational landscape of a natural aminotransferase to preferentially stabilize a less populated but reactive conformation, and thereby increase catalytic efficiency with a non-native substrate to alter substrate selectivity. Steady-state kinetics of designed variants revealed selectivity switches of up to 1900-fold, and structural analyses by room-temperature X-ray crystallography and multitemperature nuclear magnetic resonance spectroscopy confirmed that conformational equilibria favoured the target conformation. Our computational approach opens the door to the fine-tuning of enzyme conformational landscapes to create designer biocatalysts with tailored functionality.

## Main text

Enzymes are flexible macromolecules that sample multiple structural states, described by a conformational energy landscape (*1*). It is the relative stability of these conformational states, and the ability of enzymes to transition between them, that ultimately dictates enzymatic function (*2-4*). Analyses of directed evolution trajectories have shown that evolution can reshape enzyme conformational landscapes by enriching catalytically productive states and depopulating non-productive ones (*5-7*), leading to enhanced catalytic activity. Similar mechanisms contribute to the evolution of substrate selectivity, where transient conformations responsible for activity on non-cognate substrates become enriched (*8-10*). Thus, it should be possible to harness the pre-existing conformational plasticity of enzymes to rationally design biocatalysts with tailored catalytic properties by fine-tuning their conformational landscapes. However, predicting the effect of mutations on these energy landscapes remains challenging, and current computational enzyme design protocols, which focus on a single structural state, are poorly optimized for this task. New design methodologies are therefore required for the targeted alteration of subtle conformational states and equilibria, which would in turn facilitate the design of biocatalysts with customized activity and selectivity.

Here, we report a computational procedure for rationally tuning enzyme conformational landscapes that is based on multistate computational protein design, a methodology that allows protein sequences to be optimized on multiple structural states (*11*). As a case study, we remodelled the conformational landscape of aspartate aminotransferase, an enzyme that switches between open and closed conformations via hinge movement, which involves the rotation of a protein domain relative to another around an axis between two planes. Using our approach, we enriched the less populated but catalytically active closed conformation in order to increase catalytic efficiency (*k*_cat_/*K*_M_) with the non-native substrate l-phenylalanine, leading to altered substrate selectivity. Steady-state kinetics revealed *k*_cat_/*K*_M_ increases of up to 100-fold towards this aromatic amino acid, resulting in a selectivity switch of up to 1900-fold, and structural analyses by room-temperature X-ray crystallography and multitemperature nuclear magnetic resonance (NMR) spectroscopy confirmed that the conformational landscape was remodelled to favour the target state. Our methodology for conformational landscape fine-tuning could be incorporated into *de novo* enzyme design pipelines, opening the door to the creation of more complex and efficient designer biocatalysts than previously possible.

## Results

### Computational remodelling of conformational landscape

*E. coli* aspartate aminotransferase (AAT) is a pyridoxal phosphate (PLP)-dependent enzyme that catalyses the reversible transamination of l-aspartate with α-ketoglutarate, yielding oxaloacetate and L-glutamate (Supplementary Figure 1a). During its catalytic cycle, AAT undergoes hinge movement to switch between an open conformation, in the ligand-free form, and a closed conformation upon association with substrates or inhibitors (Figure 1a,b) (*12*). Previously, a hexamutant (HEX) of AAT that is approximately two orders of magnitude more catalytically efficient than the wild type (WT) with the aromatic amino acid l-phenylalanine (Table 1, Supplementary Figure 1b), and similarly efficient for transamination of l-aspartate, was engineered by replacing six of the 19 residues that are strictly conserved in AAT enzymes by those found at corresponding positions in the homologous *E. coli* tyrosine aminotransferase (*13*). Unexpectedly, it was found that unlike WT, HEX was closed in its ligand-free form (Figure 1c) (*14*), demonstrating that those six mutations shifted its conformational equilibrium to favour the closed state (Figure 1d), which is the active conformation of the enzyme (*15*). This equilibrium shift was accompanied by enhanced l-phenylalanine transamination activity resulting from a larger increase in affinity for this non-native substrate than for the native l-aspartate substrate (*13*). Thus, we postulated that we could rationally remodel the conformational landscape of AAT by using multistate computational protein design (*11*) to identify novel mutation combinations that can preferentially stabilize the closed conformation over the open conformation, and in doing so, increase substrate selectivity for l-phenylalanine.

**Figure 1.**
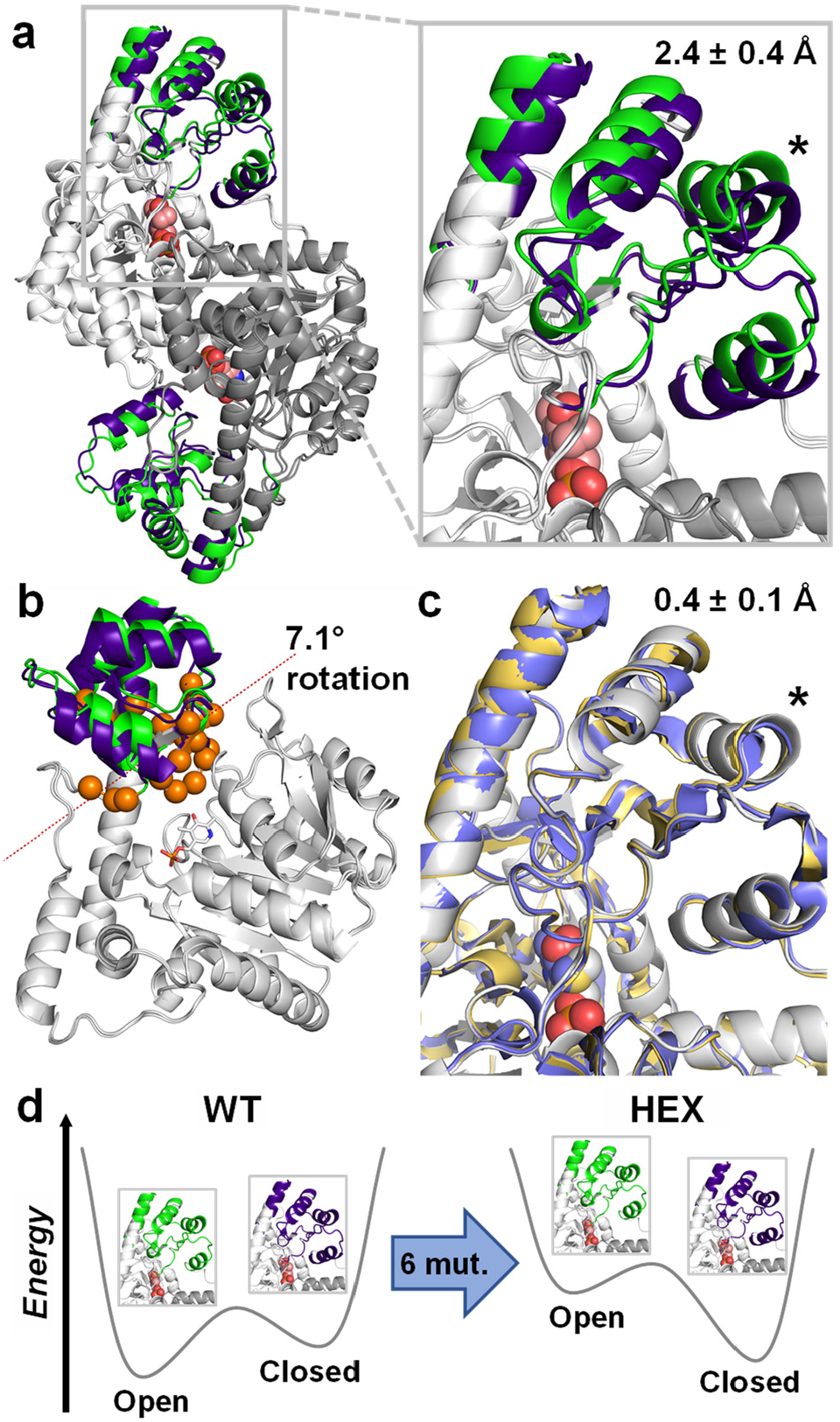
AAT conformational landscape. (a) *E. coli* AAT is a 90 kDa homodimer that undergoes a conformational change from an open (green, PDB ID: 1ARS) to closed (dark blue, PDB ID: 1ART) state upon substrate binding. This conformational transition involves rotation of a small moving domain (colored) relative to a fixed domain (white and grey for chains A and B, respectively), which causes a 2.4 ± 0.4 Å displacement (mean C_α_ distance ± s.d.) of the helix formed by residues K355–F365 (indicated by an asterisk). The PLP cofactor bound at the active site is shown as spheres (salmon). (b) Hinge movement analysis of chain A reveals a 7.1-degree rotation of the moving domain relative to the fixed domain along an axis between two planes (dotted line). Hinge-bending residues and PLP are shown as orange spheres and white sticks, respectively. (c) Superposition of HEX structures in the absence (yellow, PDB ID: 1AHE) and presence (blue, PDB ID: 1AHY) of bound inhibitor with that of the WT closed state (white, PDB ID: 1ART) show that this mutant is closed in both cases. C_α_ displacement (mean ± s.d.) of residues K355–F365 (asterisk) is indicated. (d) These results demonstrate that the six mutations (mut.) of HEX remodel its conformational landscape to favour the closed conformation.

**Table 1.** Apparent kinetic parameters of *E. coli* AAT and its mutants for transamination of various amino-acid donors with α-ketoglutarate as acceptor

| Enzyme<sup>a</sup> | Mutations |  |  |  | L-Aspartate<sup>b</sup> |  |  | L-Phenylalanine<sup>b</sup> |  |  | Selectivity<sup>c</sup> |
| --- | --- | --- | --- | --- | --- | --- | --- | --- | --- | --- | --- |
| | V35 | K37 | T43 | N64 | $K_M$<br>(mM) | $k_{cat}$<br>(s<sup>-1</sup>) | $k_{cat}/K_M$<br>(M<sup>-1</sup> s<sup>-1</sup>) | $K_M$<br>(mM) | $k_{cat}$<br>(s<sup>-1</sup>) | $k_{cat}/K_M$<br>(M<sup>-1</sup> s<sup>-1</sup>) | |
| Controls |  |  |  |  |  |  |  |  |  |  |  |
| WT | - | - | - | - | 0.21 ± 0.03 | 7.8 ± 0.3 | 37000 ± 5000 | N.D.<sup>e</sup> | N.D.<sup>e</sup> | 400 ± 100 | 0.01 |
| HEX<sup>d</sup> | L | Y | I | L | 0.059 ± 0.007 | 1.32 ± 0.03 | 22000 ± 3000 | 0.27 ± 0.03 | 9.0 ± 0.2 | 33000 ± 4000 | 1.5 |
| Closed Library |  |  |  |  |  |  |  |  |  |  |  |
| IYIT | I | Y | I | T | 0.014 ± 0.002 | 0.155 ± 0.005 | 11000 ± 2000 | 0.34 ± 0.03 | 10.1 ± 0.2 | 30000 ± 3000 | 2.7 |
| VFIT | - | F | I | T | 0.027 ± 0.003 | 0.90 ± 0.01 | 33000 ± 4000 | 0.90 ± 0.06 | 22.8 ± 0.4 | 25000 ± 2000 | 0.8 |
| VFIY | - | F | I | Y | 0.12 ± 0.01 | 0.263 ± 0.005 | 2200 ± 200 | 0.42 ± 0.04 | 17.3 ± 0.4 | 41000 ± 4000 | 19 |
| VYIT | - | Y | I | T | 0.08 ± 0.01 | 0.37 ± 0.01 | 4600 ± 600 | 1.11 ± 0.07 | 41.8 ± 0.6 | 38000 ± 2000 | 8 |
| VYIY | - | Y | I | Y | 0.09 ± 0.02 | 0.244 ± 0.006 | 2700 ± 600 | 0.58 ± 0.02 | 20.9 ± 0.2 | 36000 ± 1000 | 13 |
| Open<sub>Low</sub> Library |  |  |  |  |  |  |  |  |  |  |  |
| IFCA | I | F | C | A | 0.031 ± 0.004 | 2.63 ± 0.06 | 80000 ± 10000 | 2.3 ± 0.2 | 47 ± 1 | 20000 ± 2000 | 0.25 |
| MFCA | M | F | C | A | 0.018 ± 0.003 | 1.21 ± 0.02 | 70000 ± 10000 | 1.08 ± 0.08 | 35.9 ± 0.6 | 33000 ± 3000 | 0.47 |
| VFCA | - | F | C | A | 0.050 ± 0.006 | 4.4 ± 0.1 | 90000 ± 10000 | 4.2 ± 0.2 | 47 ± 1 | 11200 ± 600 | 0.12 |
| VFCS | - | F | C | S | 0.068 ± 0.007 | 6.0 ± 0.1 | 88000 ± 9000 | 12 ± 1 | 87 ± 3 | 7200 ± 700 | 0.08 |
| Open<sub>High</sub> Library |  |  |  |  |  |  |  |  |  |  |  |
| AIFS | A | I | F | S | 0.32 ± 0.03 | 2.42 ± 0.04 | 7600 ± 700 | 2.6 ± 0.2 | 0.54 ± 0.01 | 210 ± 20 | 0.03 |
| CIFC | C | I | F | C | 0.13 ± 0.01 | 2.62 ± 0.04 | 20000 ± 2000 | 5.9 ± 0.4 | 3.89 ± 0.08 | 660 ± 50 | 0.03 |
| CIFS | C | I | F | S | 0.28 ± 0.02 | 0.65 ± 0.01 | 2300 ± 200 | 5.4 ± 0.4 | 0.257 ± 0.006 | 48 ± 4 | 0.02 |
| SIFH | S | I | F | H | 0.33 ± 0.03 | 1.11 ± 0.02 | 3400 ± 300 | 3.7 ± 0.3 | 0.245 ± 0.008 | 66 ± 6 | 0.02 |
| SIFS | S | I | F | S | 0.35 ± 0.04 | 1.04 ± 0.02 | 3000 ± 300 | 3.9 ± 0.2 | 0.294 ± 0.006 | 75 ± 4 | 0.03 |

<sup>a</sup> Mutants are named on the basis of the amino-acid identity at the four designed positions. For example, the VFIT mutant from the Closed Library contains Val, Phe, Ile, and Thr residues at positions 35, 37, 43, and 64, respectively. WT and HEX refer to wild-type AAT and the previously published hexamutant (I3), respectively.
<sup>b</sup> All experiments were performed in triplicate using a single enzyme batch. Errors of regression fitting, which represent the absolute measure of the typical distance that each data point falls from the regression line, are provided. Concentrations of α-ketoglutarate used to determine apparent kinetic parameters of donor substrates are reported on Supplementary Table 3.
<sup>c</sup> Selectivity is defined as $(k_{cat}/K_M \text{ L-phenylalanine}) / (k_{cat}/K_M \text{ L-aspartate})$ .
<sup>d</sup> This variant also contains the T104S and N285S active-site mutations.
<sup>e</sup> Individual parameters $K_M$ and $k_{cat}$ could not be determined accurately because saturation was not possible at the maximum substrate concentration tested (40 mM, Supplementary Figure 2), which is the substrate's solubility limit. Catalytic efficiency ( $k_{cat}/K_M$ ) was therefore calculated from the slope of the linear portion ( $[S] \ll K_M$ ) of the Michaelis-Menten model ( $v_0 = (k_{cat}/K_M)[E_0][S]$ ).

To test this hypothesis, we implemented a computational strategy (Figure 2) that proceeds in five steps: (1) identification of hinge-bending residues involved in transition between open and closed conformations; (2) generation of structural ensembles approximating backbone flexibility to model open and closed conformational states; (3) optimization of side-chain rotamers for all allowed amino-acid combinations at key hinge-bending residues and neighboring positions, on each ensemble; (4) calculation of energy differences between open and closed states to predict preferred conformation, and (5) combinatorial library design using computed energy differences to select mutant sequences for experimental testing.

**Figure 2.**
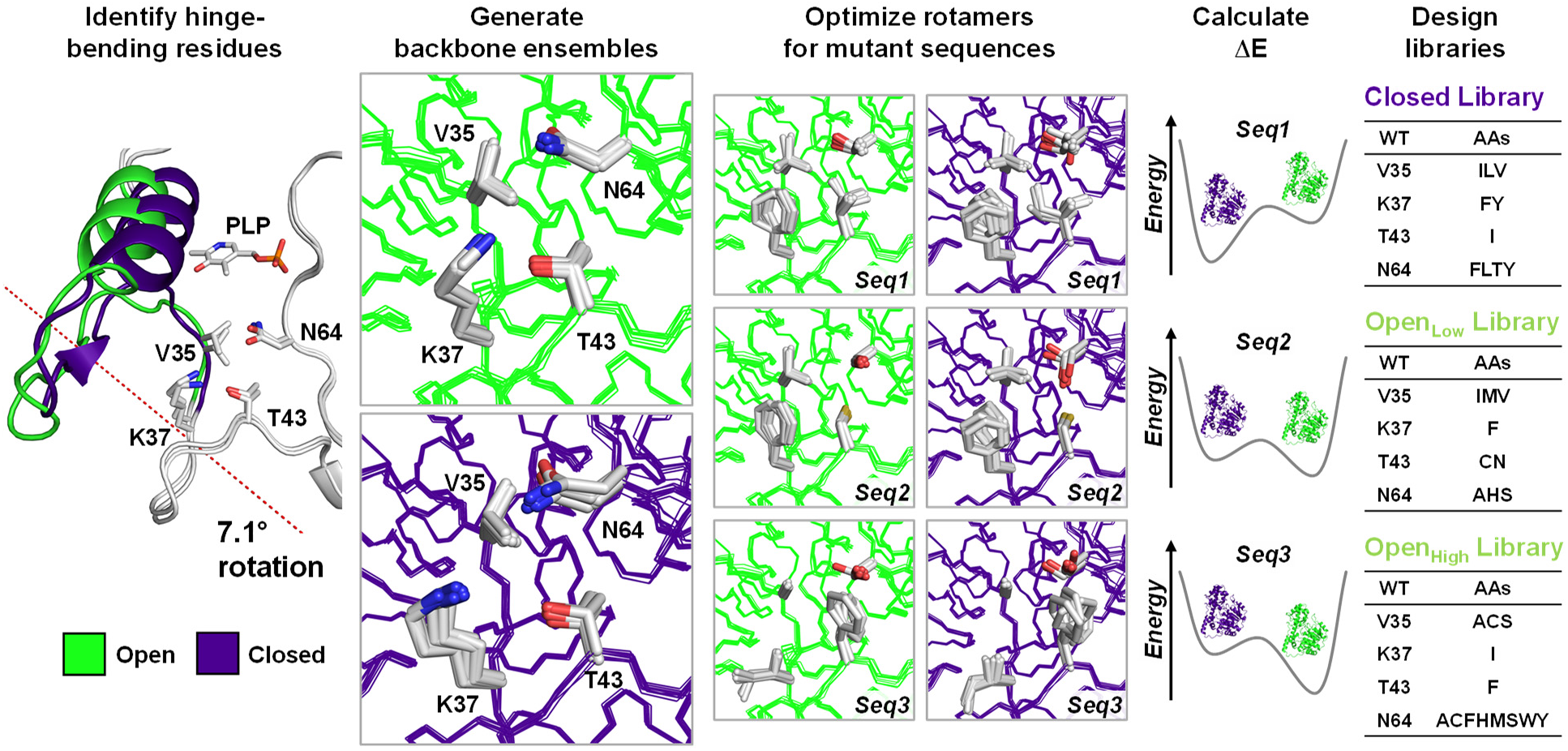
Computational remodelling of AAT conformational landscape by multistate design. To remodel the AAT conformational landscape, we followed a 5-step process: (1) identification of hinge-bending residues involved in transition between open (green) and closed (dark blue) conformational states; (2) generation of structural ensembles approximating backbone flexibility to model open and closed states; (3) optimization of rotamers for mutant sequences on both open- and closed-state ensembles; (4) calculation of energy differences between conformational states (ΔE = E_closed_ ‒ E_open_) to predict equilibrium of each mutant, and (5) combinatorial library design using ΔE values to generate Closed, Open_Low_ and Open_High_ libraries for experimental testing.

To identify hinge-bending residues for the open/closed conformational transition, crystal structures of WT AAT in its open and closed forms (PDB ID: 1ARS and 1ART, respectively (*12*)) were used as input for hinge movement analysis with DynDom, a program that identifies domains, hinge axes and hinge bending residues in proteins for which two conformations are available (*16*). DynDom analysis (Supplementary Table 1) revealed a small moving domain (Figure 1b) that rotates by 7.1 degrees about the hinge axis from the larger fixed domain, and identified 25 hinge-bending residues. We selected two of these residues, Val35 and Lys37 (numbering based on Uniprot sequence P00509), for design because they are found on the flexible loop connecting the moving and fixed domains (Figure 2). We also selected for design residues Thr43 and Asn64, which are not part of the hinge, but whose side chains form tight packing interactions with those of Val35 and Lys37. Interestingly, these residues comprise four of the six positions that were mutated in HEX (Table 1), demonstrating that our analysis using only WT structures led to the identification of positions that contribute to controlling the open/closed conformational equilibrium in AAT.

Next, we generated backbone ensembles from the open- and closed-state crystal structures to approximate the intrinsic flexibility of these two conformational states using the PertMin algorithm (*17*), which we previously showed to result in improved accuracy of protein stability predictions when used as templates in multistate design (*18*). Using the protein design software Phoenix (*19, 20*), we optimized rotamers for all combinations of proteinogenic amino acids with the exception of proline at the four designed positions on each backbone ensemble, yielding Boltzmann-weighted average energies for 130,321 (19^4^) AAT sequences that reflect their predicted stability on each conformational state. To identify mutant sequences that preferentially stabilized the closed conformation, we computed the energy difference between closed and open state ensembles (ΔE = E_closed_ – E_open_) for each sequence. As a final step, we used these ΔE values as input to the CLEARSS library design algorithm (*19*) to generate a 24-member combinatorial library of AAT mutants predicted to favour the closed state (Closed library, Supplementary Table 2) with a range of values (–86.0 to –9.5 kcal mol^−1^) encompassing that of HEX (–76.8 kcal mol^−1^, Table 2). As controls, we also generated two libraries of sequences predicted to favour the open conformation (Supplementary Table 2): the Open_Low_ library, which contains 18 sequences predicted to stabilize the open state with ΔE values (0.8–16.4 kcal mol^−1^) comparable to that of the WT (14.2 kcal mol^−1^, Table 2), and the Open_High_ library, which contains 24 sequences predicted to more strongly favour the open state due to substantial destabilization of the closed state by >120 kcal mol^−1^. While we postulated that the Open_Low_ library would yield mutants with wild-type-like conformational landscapes and therefore similar catalytic efficiency and substrate selectivity, we hypothesized that Open_High_ library mutants would be less efficient than WT with both native and non-native substrates due to their strong destabilization of the closed conformation, which is the active form of the enzyme (*15*). Thus, experimental characterization of these three mutant libraries, which comprise non-overlapping sequences (Figure 2), allowed us to assess the ability of the ΔE metric to predict sequences with conformational landscapes favouring the open or closed states.

**Table 2.** Conformational equilibrium of AAT variants

| Enzyme | Computational<sup>a</sup> |  |  | Experimental<sup>b</sup> |  |  |  |  |
| --- | --- | --- | --- | --- | --- | --- | --- | --- |
|  | E<sub>closed</sub><br>(kcal mol<sup>-1</sup>) | E<sub>open</sub><br>(kcal mol<sup>-1</sup>) | ΔE<br>(kcal mol<sup>-1</sup>) | ΔH<br>(kcal mol<sup>-1</sup>) | ΔS<br>(kcal mol<sup>-1</sup> K<sup>-1</sup>) | ΔC<sub>p</sub><br>(kcal mol<sup>-1</sup> K<sup>-1</sup>) | ΔG, 278 K<br>(kcal mol<sup>-1</sup>) | ΔG, 303 K<br>(kcal mol<sup>-1</sup>) |
| WT | -358.0 | -372.2 | 14.2 | N.D. | N.D. | N.D. | N.D. | N.D. |
| HEX | -350.2 | -273.4 | -76.8 | -9.9 | -0.032 | 0.972 | -1.72 | -0.31 |
| VFIT | -366.1 | -320.6 | -45.5 | -10.7 | -0.038 | 0.667 | -0.59 | 0.78 |
| VFIY | -357.8 | -300.0 | -57.8 | -19.6 | -0.070 | 0.369 | -0.38 | 1.59 |
| VFCS | -385.6 | -389.7 | 4.2 | N.D. | N.D. | N.D. | N.D. | N.D. |
| AIFS | -237.2 | -364.4 | 127.2 | N.D. | N.D. | N.D. | N.D. | N.D. |

<sup>a</sup> Boltzmann-weighted average potential energies (T = 300 K) for the open and closed state ensembles were computed using the Phoenix energy function (Methods). The energy difference reported corresponds to E<sub>closed</sub> - E<sub>open</sub>.
<sup>b</sup> Differences in enthalpy (ΔH), entropy (ΔS), heat capacity (ΔC<sub>p</sub>) and Gibbs free energy (ΔG) are given in the direction of enzyme closing (e.g., G<sub>closed</sub> - G<sub>open</sub>), and were calculated using a reference temperature of 298 K. N.D. indicates “not determined”.

### Kinetic analysis of designs

We screened the three mutant libraries for transamination activity with the non-native substrate l-phenylalanine (Methods) and selected the most active mutants from each library for kinetic analysis. All selected mutants catalyzed transamination of l-phenylalanine or l-aspartate with α-ketoglutarate, and displayed substrate inhibition with this acceptor substrate, as is the case for WT (Table 1, Supplementary Tables 3–4, Supplementary Figures 2–5). All Closed library mutants displayed catalytic efficiencies towards l-phenylalanine that were improved by approximately two orders of magnitude relative to WT (Table 1), comparable to HEX, and all were similarly or more active with this non-native substrate than with l-aspartate, in stark contrast with WT AAT, which prefers the native substrate by a factor of 100 (Supplementary Figure 6). By contrast, Open_Low_ and Open_High_ mutants favoured the native over the non-native substrate by up to 50-fold, similar to WT. Surprisingly, Open_Low_ mutants had lower *K*_M_ values and were more catalytically efficient than WT with both l-phenylalanine and l-aspartate, which could be due to the fact that these mutants have ΔE values similar to the WT but stabilize both the open and closed conformations by >10 kcal mol^‒1^ (Table 2, Supplementary Table 2). This is not the case for Open_High_ mutants, which have similar *K*_M_ values but are less catalytically efficient with the native substrate than WT, in agreement with our hypothesis that strong destabilization of the catalytically active closed conformation would result in less efficient catalysis. Overall, these kinetic results support the hypothesis that the change in substrate selectivity is linked to the conformational equilibrium previously suggested by the data from the HEX mutant (*13, 14*).

### Structural analysis of designs

To provide structural information on the conformations adopted by AAT mutants, we turned to room-temperature X-ray crystallography, which provides insight into enzyme conformational ensembles under conditions that are relevant to catalysis (*21*) and free of potential distortions or conformational bias introduced by sample cryocooling (*22*). We crystallized WT, HEX, and select variants from the Closed (VFIT and VFIY), Open_Low_ (VFCS) and Open_High_ (AIFS) libraries. All six enzymes yielded crystals under similar conditions (Supplementary Table 5), which could only be obtained in the presence of maleate (Supplementary Figure 1c), an inhibitor that stabilizes the closed conformation when bound by the enzyme (*23*). To obtain structures in the absence of maleate, we applied a rigorous crystal soaking method to serially dilute and extract the inhibitor from the crystallized enzymes (Methods). We collected X-ray diffraction data at room-temperature (278 K) for all variants with the exception of AIFS, which could only be measured at cryogenic temperature (100 K) because we could only obtain small crystals that were not robust to radiation damage at non-cryogenic temperatures. We applied statistical criteria (Supplementary Table 6) to assign high-resolution cut-offs of 1.37–2.31 Å for our data sets, and all structures were subsequently determined by molecular replacement in space group P63 with an enzyme homodimer in the asymmetric unit. In the inhibitor-bound state, all structures were closed as expected (Supplementary Figure 7) due to electrostatic interactions between maleate and the side chains of Arg280 and Arg374 (Supplementary Figure 8). Upon soaking crystals of the WT enzyme to remove the bound inhibitor, we observed that one subunit (chain A) within the enzyme homodimer was in the open conformation (Figure 3), confirming that the soaking procedure was able to remove the bound maleate, and that the crystal lattice could accommodate the domain rotation required for opening and closing of this subunit. In the maleate-free structures, a sulfate ion and one or more ordered water molecules occupy the inhibitor binding site (Supplementary Figure 9), as was previously observed in WT (*24*) and HEX (*14*) structures at cryogenic temperatures.

**Figure 3.**
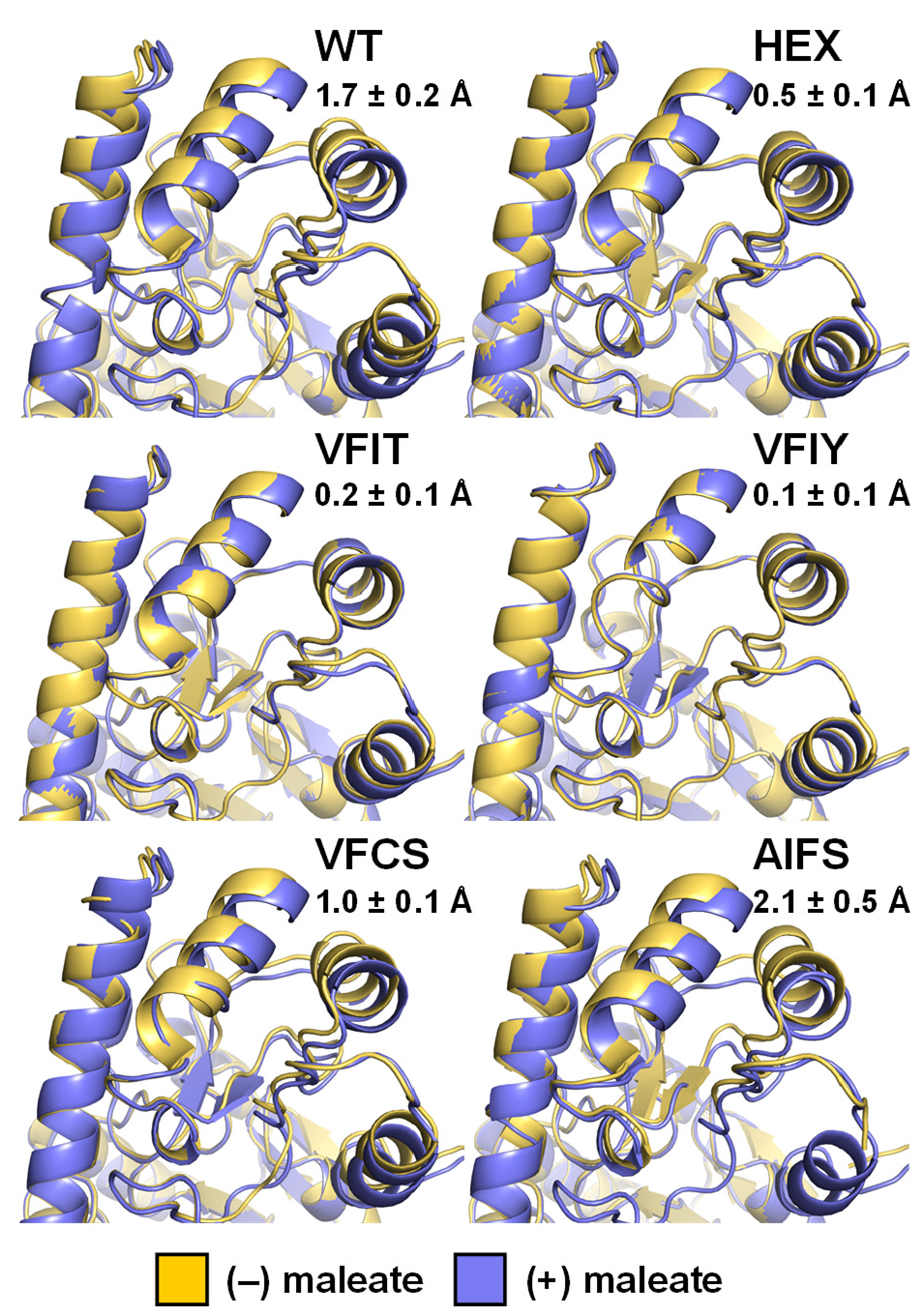
Crystal structures. Overlay of AAT structures (Chain A) in the presence and absence of maleate show transition from open to closed states upon inhibitor binding for WT, VFCS, and AIFS, but not for HEX, VFIT, and VFIY, which already adopt the closed conformation in the absence of bound inhibitor. Average displacements of helix formed by residues K355–F365 upon maleate binding are reported as the average pairwise distance of corresponding C_α_ atoms for the 11 residues comprising this helix (mean ± s.d.).

Superposition of bound and unbound structures for each variant confirmed that closed library mutants VFIT and VFIY remained in the closed conformation in the inhibitor-free form, similar to HEX, while Open_Low_ variant VFCS and Open_High_ variant AIFS adopted the open conformation, similar to WT (Figure 3). AIFS is unique in that the electron density is especially weak in the regions corresponding to the helix formed by Pro12–Leu19 and the loop connecting moving and fixed domains (Leu31–Thr43), even at cryogenic temperatures, demonstrating that these structural segments are disordered when the enzyme adopts the open conformation (Supplementary Figure 10). This result suggests that the four mutations of AIFS, three of which are found within the Leu31–Thr43 loop, contribute to destabilize the open conformation, consistent with the calculated E_open_ value of this variant (Table 2) being >7 kcal mol^−1^ higher than that of the other variants that favour the open state (WT and VFCS). Interestingly, the amplitude of the open/closed conformational transition that occurs in open variants upon maleate binding (Figure 3) correlated with their computed ΔE values (VFCS < WT < AIFS), suggesting that this metric can be used to fine-tune this enzyme conformational landscape. Furthermore, DynDom analyses of all variants confirmed that only WT, VFCS, and AIFS undergo hinge motion upon maleate binding (Supplementary Table 7), which rotates the moving domain relative to the fixed domain by 4.6, 2.6, and 5.9 degrees, respectively.

### NMR analysis of conformational landscapes

Having demonstrated crystallographically that our designed variants adopted the target conformation in the absence of ligand, we turned to NMR spectroscopy to gain insights into their conformational equilibria in solution, and compared results against those obtained for WT and HEX. We first measured ^1^H-^15^N HSQC spectra for WT in the presence and absence of l-aspartate (Supplementary Figure 11a). As expected for a protein of this size, the spectrum consisted of a large number of peaks that were broad with a high degree of overlap. It was nonetheless possible to assign the unique chemical shifts of peaks from the indole NH group for three native Trp residues by comparison of these spectra with those acquired with single Trp mutants (Supplementary Figure 11b,c). This allowed assignment of a peak that was significantly broadened in spectra of both HEX and Closed library mutant VFIY to the indole NH from Trp307 (Supplementary Figure 11d), whose side chain is closest to the designed hinge residues. Moreover, the Trp307 indole peak was no longer detectable when the l-aspartate substrate was present for both WT and mutant enzymes, most likely being broadened beyond detection. Since all conditions that favour the closed state (i.e., HEX and Closed library mutations and/or l-aspartate binding) broaden the Trp307 indole resonance, this exchange appears to be associated with the closed state, potentially due to conformational dynamics around the hinge.

In order to characterize the thermodynamics of the exchange processes around the hinge region, we labelled AAT variants at a single site with ^19^F using site-specific incorporation of the noncanonical amino acid 4-trifluoromethyl-l-phenylalanine (*25*). Phe217 was chosen as the incorporation site since it is proximal to hinge residues but is not in direct contact with the substrate (Figure 4a). The ^19^F spectrum of WT at 278 K showed a single peak centred at approximately ‒60 ppm that shifts downfield as temperature is increased (Figure 4b), similar to what is observed for free 4-trifluoromethyl-l-phenylalanine in solution (Supplementary Figure 12). By contrast, the HEX spectrum at 278 K showed a large broad peak centred at around ‒61.6 ppm, with another peak of substantially lower intensity also appearing at a similar shift to that observed in the WT spectrum (‒60.4 ppm). The relative intensity of these 2 peaks changed as the temperature was increased, with the low intensity peak increasing as the major peak decreased. This is characteristic of two-state exchange, with the equilibrium between the two states being shifted by the temperature change. Given that the crystal structure of HEX in its ligand-free form showed a closed conformation (Figure 3), it is likely that the major peak reflects a local chemical environment created by the closed state, with a small population in an open state similar to that seen in the WT spectrum.

**Figure 4.**
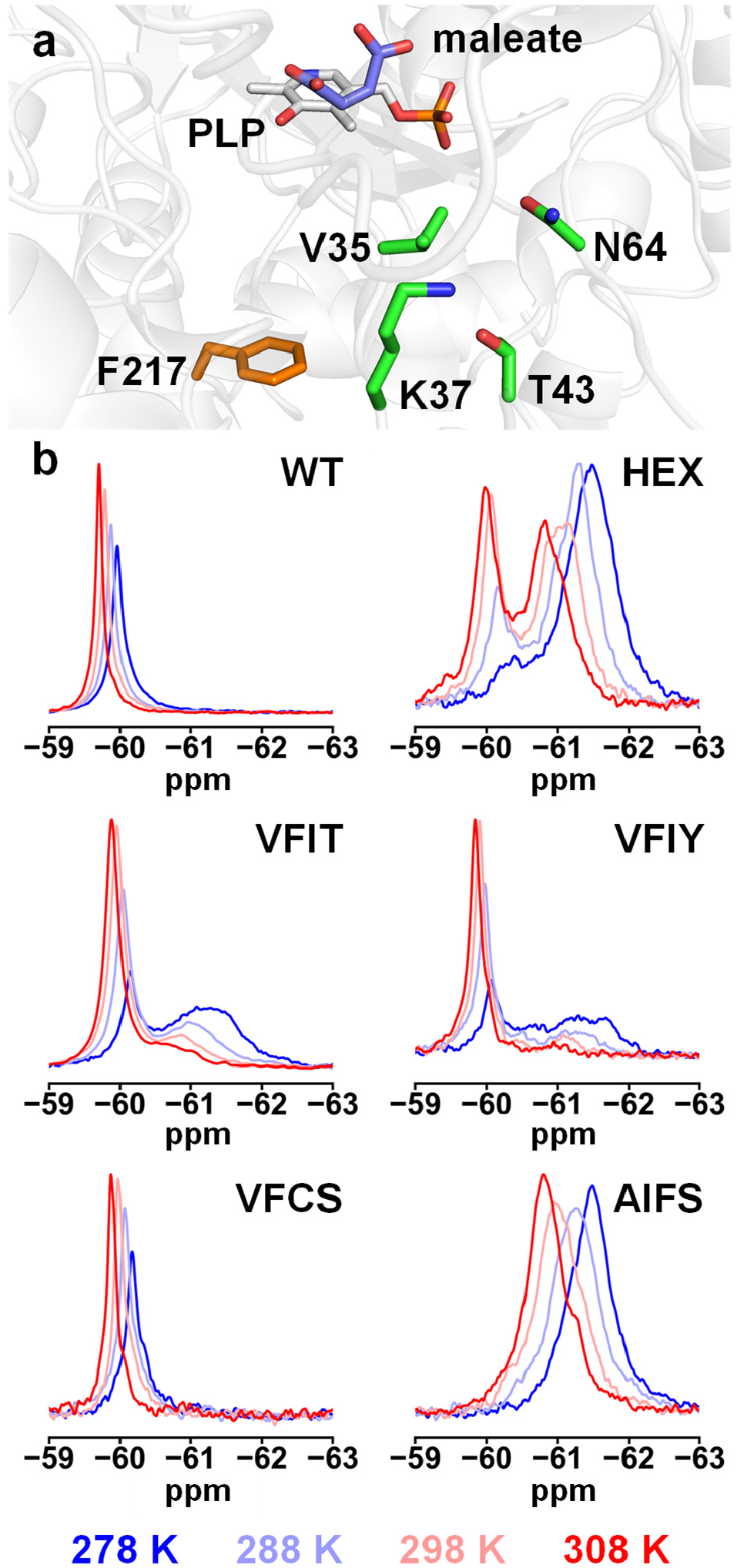
Conformational landscape analysis by NMR. (a) To evaluate conformational equilibrium of AAT variants, we introduced the fluorinated amino acid 4-trifluoromethyl-l-phenylalanine at position F217 (orange), which is located closer to hinge-bending and designed residues (green) than to the bound maleate inhibitor (blue). Crystal structure shown is that of wild-type (WT) AAT at 278 K (PDB ID: 8E9K). (b) ^19^F NMR spectra of AAT variants in the absence of ligand show dynamic equilibrium between 278 K and 308 K for HEX, VFIT, VFIY, and AIFS, confirming that these proteins are undergoing exchange. This is not the case for WT and Open_Low_ library mutant VFCS, who both adopt predominantly the open conformation within this temperature range. For HEX and Closed library mutants VFIT and VFIY, the open conformation is enriched as temperature increases. For AIFS, spectra suggest that this Open_High_ library mutant samples conformations distinct from those sampled by the other variants.

Using peak deconvolution and integration, it was possible to calculate relative populations for each species for variants undergoing the observed two-state exchange, along with the free energy difference between states (Supplementary Figure 13, Supplementary Table 8). We calculated ΔG at 278 K for HEX to be –1.72 kcal mol^−1^ (Table 2), a small difference that would be compatible with the interconversion between these two states to be part of the catalytic cycle (*26*). These peak volumes could also be calculated over the entire temperature range tested, giving rise to a van’t Hoff relationship with a small degree of curvature (Supplementary Figure 14). Deviations from linearity can occur when there are differences in heat capacity between the two states, as would be expected for a process involving a change in conformational states over the temperature range tested (*27*). By contrast, the presence of a single peak in WT spectra suggests that the closed state is not significantly populated under these conditions, as expected from its open-state crystal structures obtained under both cryogenic (*12, 24*) and room-temperature conditions.

We next analyzed two mutants from the Closed library (VFIT and VFIY). ^19^F NMR spectra at 278 K for both mutants showed a narrow peak centred at approximately ‒60.2 ppm that is similar to that of WT (Figure 4). Spectra of these variants also showed another broad peak centred at approximately ‒61.2 ppm that is similar to the HEX peak characteristic of the closed conformation. The relative intensity of the two peaks showed similar temperature dependence, with an increase in the relative intensity of the WT-like peak as temperature was increased to 308 K. Peak deconvolution and integration (Supplementary Figure 13–14, Supplementary Table 8) were used to evaluate the population of the two states, and confirmed that at low temperatures both VFIT and VFIY favour the state resembling that adopted by the closed HEX mutant (T < 288 K or 283 K, respectively). However, unlike HEX, at higher temperatures the alternate conformation with the WT-like peak becomes the favoured state (Table 2). Interestingly, ΔE values for these mutants were smaller in magnitude than that calculated for HEX, supporting the predictive nature of the calculated energy differences between open and closed states for these sequences.

To determine if these differences in the temperature dependence of exchange could be observed in the crystal state, we also solved the maleate-free structures of WT, HEX, and VFIT at 303 K (Supplementary Table 9, Supplementary Figure 9), and calculated isomorphous difference density maps by subtracting electron density at 278 K (Supplementary Figure 15). Comparing

VFIT data obtained at 278 K and 303 K results in substantial difference density throughout chain A, the chain that opens when WT crystals are soaked to remove maleate. By contrast, similar comparisons for WT and HEX showed relatively little difference density. This analysis confirms that VFIT undergoes larger local conformational changes than either HEX or WT when the temperature of the crystal is increased to 303 K, in agreement with our van’t Hoff analysis of NMR data (Table 2, Supplementary Table 8). The agreement between temperature-dependent X-ray crystallography and NMR data provides strong evidence that the conformational exchange detected in the NMR experiments reflects a dynamic equilibrium between closed and open conformations, with mutations that favour the closed conformation also increasing selectivity toward l-phenylalanine.

We next analyzed two mutants designed to favour the open conformation in their ligand-free forms. Open_Low_ mutant VFCS showed ^19^F NMR spectra that were very similar to those of WT within the tested temperature range (Figure 4b, Supplementary Figure 13), consistent with its preference for the open conformation (Figure 3, Table 2). However, Open_High_ mutant AIFS gave rise to spectra that were distinct from those of all other variants (Figure 4b), but could be deconvoluted to two exchanging peaks (Supplementary Figure 13), to allow estimation of populations (Supplementary Table 8, Supplementary Figure 14). We postulate that those peaks correspond to alternate open conformations distinct from the one sampled by the other variants. This hypothesis is supported by our observations that ligand-free AIFS is open at low temperature (Figure 3) but contains disordered segments around the Pro12–Leu19 helix and the loop containing three of the four designed positions (Supplementary Figure 10), which are located close to the Phe217 position where the ^19^F label was introduced. The observed heterogeneity could therefore correspond to a mixture of these alternate open conformations. Additional support that the exchange detected in AIFS differed from that of HEX and closed mutants was provided by the van’t Hoff analysis, which showed no curvature for AIFS unlike for the closed mutants (Supplementary Figure 14), suggesting that the Open_High_ variant does not undergo the open/closed conformational transition within this temperature range.

## Discussion

Here, we successfully remodeled the conformational landscape of an enzyme via targeted alterations to the equilibrium between two distinct conformational states related by a hinge-bending motion. The resulting equilibrium shift promoted activity towards a non-native substrate, leading to a selectivity switch of up to 1900-fold. As many enzymes undergo hinge-mediated domain motions during their catalytic cycles (*28*), the multistate design approach presented here should be straightforward to implement for such enzymes. Given the ability of our design procedure to distinguish between closed and open states whose free energy difference is on the order of a single hydrogen bond, our approach could, in principle, also be applied to preferentially stabilize catalytically competent substates involving more subtle structural changes, such as backbone carbonyl flips (*29*) or side-chain rotations (*30*). This methodology could therefore help to tailor catalytic efficiency or substrate selectivity by mimicking, *in silico*, the processes of evolution that harness altered conformational equilibria to tune function (*7, 31, 32*).

The predictive capacity of our multistate design framework could only be achieved by evaluating the energy of sequences on multiple conformational states. For example, the VFCS variant that prefers the open state is predicted to be more stable on the closed state than Closed library mutants VFIT and VFIY (Table 2), and more stable on the Open state than AIFS even though it is less open than this variant (Figure 3). Furthermore, there were no obvious trends in designed mutations that could explain their effect on the conformational landscape, as none of these introduced bulkier or smaller amino acids at all or specific residue positions to cause or alleviate steric clashes in the open or closed conformations so as to shift the equilibrium towards one of these states, which has been the approach others have used to shift conformational equilibria (*33*). Thus, subtle effects of mutation combinations on the relative stability of each conformational state were likely responsible for the observed preference of mutants for the open or closed conformations.

Our results demonstrate the utility of multistate design, with ΔE values calculated from ensemble energies of open and closed states, for the targeted alteration of subtle conformational equilibria, an approach that represents a useful alternative to heuristic methods that others have used to tune the relative stability of protein conformational states (*34*). Extending this concept, we envision that de novo design of artificial enzymes with native-like catalytic efficiency and selectivity for complex multistep chemical transformations will require a holistic approach where every conformational state and/or substate required to stabilize reaction intermediates and transition states are explicitly modelled, and their relative energies optimized. The multistate design method for conformational landscape remodelling presented here could therefore be incorporated into *de novo* enzyme design pipelines, helping to bridge the gap between the carving of an active site for transition-state stabilization, and the modulation of conformational dynamics required for efficient passage along the reaction coordinate.

## Supporting information

Restraints for PLP refinement

