## Supplementary Information for "Computational remodeling of an enzyme conformational landscape for altered substrate selectivity"

**This file includes:**

Supplementary Tables 1–10

Supplementary Figures 1–15

Supplementary Methods

Supplementary References

**Supplementary Table 1**. Hinge movement analysis for wild–type *E. coli* AAT

| DynDom Results | 1ARS *vs.* 1ART [a] |
|---|---|
| **Fixed Domain Residues** | 31, 36–324, 339–342, 345–348, 375 |
| **Moving Domain Residues** | 13–30, 32–35, 325–338, 343–344, 349–374, 376–394 |
| **Unassigned Residues** | 1–12, 395–396 |
| **Angle of rotation (°)** | 7.1 |
| **Translation along axis (Å)** | –0.4 |
| **Closure (%)** | 100.0 |
| **Bending Residues** | 29–32, 35–37, 323–325, 338–349, 374–376 |

[a] Amino-acid residues are numbered according to the AAT sequence (Uniprot ID: P00509). The 1ARS and 1ART crystal structures of wild-type *E. coli* AAT were used to represent the open and closed states, respectively. For both structures, unit cells of similar dimensions (1ARS: a = 155.42 Å, b = 87.13 Å, c = 79.4 Å; 1ART: a = 157.14 Å, b = 85.51 Å, c = 78.93 Å) corresponding to space group C 2 2 $2_1$ contain a single protein chain. All analyses performed using DynDom (*35*).

**Supplementary Table 2.** Computed energies of individual mutant sequences from each designed combinatorial library

| **Closed Library** [a] | | | | **$\text{Open}_{\text{Low}}$ Library** [a] | | | | **$\text{Open}_{\text{High}}$ Library** [a] | | | |
|---|---|---|---|---|---|---|---|---|---|---|---|
| **Mutant** [b] | **$E_{closed}$** [c] **(kcal mol$^{-1}$)** | **$E_{open}$** [c] **(kcal mol$^{-1}$)** | **ΔE** [c] **(kcal mol$^{-1}$)** | **Mutant** [b] | **$E_{closed}$** [c] **(kcal mol$^{-1}$)** | **$E_{open}$** [c] **(kcal mol$^{-1}$)** | **ΔE** [c] **(kcal mol$^{-1}$)** | **Mutant** [b] | **$E_{closed}$** [c] **(kcal mol$^{-1}$)** | **$E_{open}$** [c] **(kcal mol$^{-1}$)** | **ΔE** [c] **(kcal mol$^{-1}$)** |
| IFIF | –358.9 | –272.9 | –86.0 | IFNH | –387.7 | –388.5 | 0.8 | AIFF | –231.5 | –359.9 | 128.4 |
| LFIL | –354.6 | –277.2 | –77.4 | IFCH | –384.1 | –385.1 | 1.1 | AIFW | –231.1 | –359.4 | 128.2 |
| LYIL | –350.2 | –273.4 | –76.8 | VFNS | –390.6 | –391.8 | 1.2 | CIFM | –239.9 | –367.5 | 127.6 |
| IFIL | –353.3 | –280.0 | –73.3 | VFNA | –386.8 | –388.3 | 1.5 | AIFY | –231.2 | –358.8 | 127.6 |
| IFIY | –357.3 | –291.6 | –65.7 | VFNH | –388.6 | –391.6 | 3.0 | AIFS | –237.2 | –364.4 | 127.2 |
| VFIY | –357.8 | –300.0 | –57.8 | IFNA | –385.2 | –388.6 | 3.4 | AIFM | –235.3 | –362.4 | 127.1 |
| VFIL | –356.6 | –298.8 | –57.8 | IFNS | –388.5 | –392.1 | 3.7 | CIFS | –241.7 | –368.5 | 126.9 |
| IYIF | –306.9 | –250.2 | –56.7 | VFCS | –385.6 | –389.7 | 4.2 | AIFH | –239.3 | –365.5 | 126.1 |
| VYIL | –350.7 | –295.4 | –55.3 | VFCA | –381.7 | –386.3 | 4.6 | SIFF | –232.8 | –358.6 | 125.9 |
| VYIY | –349.5 | –294.6 | –54.9 | VFCH | –384.8 | –389.5 | 4.7 | SIFS | –238.1 | –363.8 | 125.7 |
| VYIF | –352.8 | –302.4 | –50.4 | IFCA | –381.3 | –387.8 | 6.5 | CIFH | –244.1 | –369.4 | 125.3 |
| IFIT | –361.7 | –313.4 | –48.3 | IFCS | –384.5 | –391.2 | 6.7 | SIFY | –232.1 | –357.4 | 125.3 |
| IYIL | –301.5 | –253.4 | –48.2 | MFNA | –378.8 | –390.3 | 11.4 | SIFM | –236.8 | –362.0 | 125.2 |
| VFIF | –361.9 | –315.7 | –46.2 | MFNS | –382.8 | –394.5 | 11.7 | AIFC | –235.2 | –360.1 | 124.9 |
| VFIT | –366.1 | –320.6 | –45.5 | MFCH | –368.5 | –383.9 | 15.3 | AIFA | –237.4 | –362.1 | 124.7 |
| VYIT | –355.8 | –315.2 | –40.6 | MFCS | –379.1 | –394.6 | 15.5 | CIFC | –240.0 | –364.4 | 124.3 |
| IYIY | –304.5 | –273.2 | –31.2 | MFCA | –375.0 | –390.8 | 15.7 | CIFA | –242.1 | –366.3 | 124.2 |
| LFIF | –321.3 | –297.2 | –24.2 | MFNH | –371.4 | –387.8 | 16.4 | CIFW | –238.7 | –362.8 | 124.1 |
| LYIF | –316.6 | –295.2 | –21.4 | | | | | SIFH | –240.3 | –364.1 | 123.9 |
| LFIY | –334.7 | –320.3 | –14.4 | | | | | SIFC | –236.0 | –359.3 | 123.3 |
| LFIT | –353.9 | –341.9 | –12.0 | | | | | SIFA | –238.3 | –361.5 | 123.2 |
| LYIY | –329.8 | –318.3 | –11.5 | | | | | CIFF | –236.1 | –358.3 | 122.2 |
| IYIT | –306.4 | –294.9 | –11.5 | | | | | SIFW | –236.2 | –358.3 | 122.1 |
| LYIT | –348.9 | –339.5 | –9.5 | | | | | CIFY | –236.2 | –356.9 | 120.7 |

[a] Closed, $\text{Open}_{\text{Low}}$, and $\text{Open}_{\text{High}}$ libraries comprise 24, 18, and 24 combinatorial mutant sequences, respectively.
[b] Mutants are named on the basis of the amino-acid identity at designed positions. For example, the VFIY mutant from the Closed library contains Val, Phe, Ile, and Tyr residues at positions 35, 37, 43, and 64, respectively.
[c] Boltzmann-weighted average potential energies (T = 300 K) for the closed ($E_{closed}$) and open ($E_{open}$) state ensembles were computed using the Phoenix energy function (Methods). The energy difference (ΔE) corresponds to $E_{closed} – E_{open}$.

**Supplementary Table 3.** Acceptor substrate concentrations used to measure apparent kinetic parameters reported on Table 1

| Enzyme | [α-ketoglutarate] (mM) |
|---|---|
| **WT** | 0.625 |
| **HEX** | 0.625 |
| **IYIT** | 0.3125 |
| **VFIT** | 0.3125 |
| **VFIY** | 0.3125 |
| **VYIT** | 0.3125 |
| **VYIY** | 0.3125 |
| **IFCA** | 0.1563 |
| **MFCA** | 0.1563 |
| **VFCA** | 0.1563 |
| **VFCS** | 0.3125 |
| **AIFS** | 1.0 |
| **CIFC** | 0.625 |
| **CIFS** | 1.25 |
| **SIFH** | 1.25 |
| **SIFS** | 1.25 |

**Supplementary Table 4.** Apparent kinetic parameters of *E. coli* AAT and its mutants for transamination of α-ketoglutarate as an acceptor with various amino-acid donors

| Enzyme | α-ketoglutarate [a] | | | |
|---|---|---|---|---|
| | $K_M$ (mM) | $K_I$ (mM) | $k_{cat}$ ($s^{-1}$) | $k_{cat}/K_M$ ($M^{-1}$ $s^{-1}$) |
| **Controls** | | | | |
| **WT** [b] | 0.065 ± 0.005 | 4.0 ± 0.3 | 12.2 ± 0.3 | 190000 ± 20000 |
| **HEX** [b] | 0.028 ± 0.003 | 2.8 ± 0.3 | 1.83 ± 0.06 | 65000 ± 7000 |
| **Closed Library** | | | | |
| **IYIT** [c] | 0.26 ± 0.06 | 1.7 ± 0.4 | 16 ± 2 | 60000 ± 20000 |
| **VFIT** [b] | 0.050 ± 0.004 | 2.7 ± 0.2 | 1.15 ± 0.03 | 23000 ± 2000 |
| **VFIY** [c] | 0.09 ± 0.02 | 1.6 ± 0.3 | 20 ± 2 | 220000 ± 50000 |
| **VYIT** [c] | 0.10 ± 0.02 | 1.3 ± 0.2 | 43 ± 3 | 430000 ± 90000 |
| **VYIY** [c] | 0.049 ± 0.007 | 1.6 ± 0.2 | 26 ± 1 | 530000 ± 80000 |
| **$Open_{Low}$ Library** | | | | |
| **IFCA** [c] | 0.017 ± 0.001 | 2.0 ± 0.2 | 53 ± 1 | 3100000 ± 200000 |
| **MFCA** [c] | 0.0128 ± 0.0008 | 2.4 ± 0.2 | 46.3 ± 0.8 | 3600000 ± 200000 |
| **VFCA** [c] | 0.019 ± 0.001 | 1.8 ± 0.2 | 56 ± 1 | 2900000 ± 200000 |
| **VFCS** [b] | 0.041 ± 0.006 | 2.6 ± 0.3 | 7.9 ± 0.4 | 190000 ± 30000 |
| **$Open_{High}$ Library** | | | | |
| **AIFS** [b] | 0.17 ± 0.01 | 4.3 ± 0.3 | 3.6 ± 0.1 | 21000 ± 1000 |
| **CIFC** [d] | 0.10 ± 0.01 | 4.8 ± 0.6 | 2.6 ± 0.1 | 26000 ± 3000 |
| **CIFS** [d] | 0.35 ± 0.04 | 10 ± 1 | 0.63 ± 0.03 | 1800 ± 200 |
| **SIFH** [d] | 0.14 ± 0.02 | 13 ± 3 | 0.69 ± 0.03 | 4900 ± 700 |
| **SIFS** [d] | 0.14 ± 0.01 | 14 ± 2 | 0.67 ± 0.02 | 4800 ± 500 |

[a]All experiments were performed in triplicate using a single enzyme batch. Fitting of the kinetic data was done with a rate equation that takes into account substrate inhibition by α-ketoglutarate: $v_0 = (v_{max}[S])/(K_M + [S] + [S]^2/K_I)$. Errors of regression fitting, which represent the absolute measure of the typical distance that each data point falls from the regression line, are provided.
[b] 20 mM of L-aspartate as donor substrate was used to determine apparent kinetic parameters of the α-ketoglutarate acceptor.
[c] 20 mM of L-phenylalanine as donor substrate was used to determine apparent kinetic parameters of the α-ketoglutarate acceptor.
[d] 10 mM of L-aspartate as donor substrate was used to determine apparent kinetic parameters of the α-ketoglutarate acceptor.

**Supplementary Table 5.** Crystallization conditions

| Enzyme [a] | Protein (mg mL$^{-1}$) | $(NH_4)_2SO_4$ (M) |
|---|---|---|
| **WT** | 5 | 1.79 |
| **HEX** | 10 | 1.79 |
| **VFIT** | 5 | 1.89 |
| **VFIY** | 10 | 1.79 |
| **VFCS** | 15 | 1.79 |
| **AIFS** | 10 | 1.79 |

[a] All enzymes were crystallized in a mother liquor containing 100 mM HEPES buffer (pH 7.5), 2% PEG-400, and 20 mM maleate, with varying concentrations of ammonium sulfate as noted. Protein solutions were diluted to the above concentrations with 20 mM potassium phosphate buffer (pH 7.5) containing 2 mM EDTA and 10 μM pyridoxal 5'-phosphate.

**Supplementary Table 6.** Crystallographic data and refinement statistics for structures at 100 and 278 K

| | WT | WT | HEX | HEX | VFIT | VFIT | VFIY | VFIY | VFCS | VFCS | AIFS | AIFS |
|---|---|---|---|---|---|---|---|---|---|---|---|---|
| **Maleate** | (–) | (+) | (–) | (+) | (–) | (+) | (–) | (+) | (–) | (+) | (–) | (+) |
| **PDB ID** | 8E9P | 8E9K | 8E9J | 8E9Q | 8E9L | 8E9M | 8E9N | 8E9O | 8E9R | 8E9S | 8E9C | 8E9D |
| **Data collection** [a] | | | | | | | | | | | | |
| **Temp. (K)** | 278 | 278 | 278 | 278 | 278 | 278 | 278 | 278 | 278 | 278 | 100 | 100 |
| **Resolution (Å)** | 62.51–2.09 | 62.16–1.83 | 124.48–2.09 | 81.70–1.80 | 124.38–2.31 | 124.30–1.76 | 124.43–1.88 | 124.31–1.96 | 124.46–1.90 | 124.25–2.00 | 67.54–2.18 | 61.15–1.37 |
| **Space group** | P 63 | P 63 | P 63 | P 63 | P 63 | P 63 | P 63 | P 63 | P 63 | P 63 | P 63 | P 63 |
| ***Cell params.*** | | | | | | | | | | | | |
| **a b c (Å)** | 144.33 144.33 81.12 | 143.55 143.55 81.45 | 143.74 143.74 81.65 | 143.39 143.39 81.70 | 143.62 143.62 81.66 | 143.53 143.53 81.65 | 143.67 143.67 81.53 | 143.52 143.52 81.53 | 143.63 143.63 81.41 | 143.47 143.47 81.68 | 141.63 141.63 80.92 | 141.22 141.22 81.24 |
| **α β γ (°)** | 90 90 120 | 90 90 120 | 90 90 120 | 90 90 120 | 90 90 120 | 90 90 120 | 90 90 120 | 90 90 120 | 90 90 120 | 90 90 120 | 90 90 120 | 90 90 120 |
| **Chains per asymm. unit** | 2 | 2 | 2 | 2 | 2 | 2 | 2 | 2 | 2 | 2 | 2 | 2 |
| **$R_{pim}$** | 0.077 (1.119) | 0.073 (0.955) | 0.103 (1.096) | 0.071 (1.037) | 0.182 (1.583) | 0.068 (1.076) | 0.069 (1.259) | 0.092 (0.951) | 0.070 (1.017) | 0.112 (0.722) | 0.059 (0.575) | 0.014 (0.399) |
| **$CC_{1/2}$** | 0.995 (0.227) | 0.996 (0.348) | 0.993 (0.273) | 0.994 (0.300) | 0.982 (0.295) | 0.997 (0.267) | 0.997 (0.301) | 0.993 (0.309) | 0.992 (0.303) | 0.993 (0.571) | 0.997 (0.470) | 1.000 (0.590) |
| **I/σI** | 6.8 (0.4) | 6.5 (0.5) | 5.9 (0.6) | 7.4 (0.8) | 3.3 (0.5) | 7.4 (0.6) | 7.3 (0.5) | 6.0 (0.6) | 6.1 (0.5) | 5.1 (0.7) | 8.3 (1.2) | 24.9 (1.7) |
| **Complete. (%)** | 100.0 (100.0) | 100.0 (100.0) | 99.8 (99.7) | 99.8 (99.3) | 100.0 (100.0) | 100.0 (100.0) | 98.7 (97.6) | 100.0 (99.4) | 99.8 (98.7) | 100.0 (99.1) | 100.0 (100.0) | 97.7 (92.2) |
| **Multiplicity** | 10.5 (9.3) | 10.5 (10.7) | 10.2 (10.1) | 5.0 (4.8) | 10.3 (10.3) | 10.0 (9.0) | 10.3 (10.5) | 10.1 (10.2) | 81.5 (82.7) | 40.6 (40.5) | 40.6 (38.6) | 19.8 (17.3) |
| **Wilson B-factor ($Å^2$)** | 31.230 | 23.930 | 28.090 | 23.300 | 27.750 | 21.810 | 26.890 | 23.820 | 23.940 | 21.990 | 35.942 | 16.485 |
| **# unique reflections** | 57093 (2848) | 84225 (4169) | 56805 (2782) | 88385 (4395) | 42181 (2069) | 94818 (4736) | 76900 (3787) | 68674 (3426) | 75209 (3696) | 64724 (3199) | 48326 (2387) | 188377 (8826) |
| **Refinement** | | | | | | | | | | | | |
| **R work/free** | 0.1686/ 0.2058 | 0.1528/ 0.1832 | 0.1686/ 0.1970 | 0.1435/ 0.1721 | 0.1829/ 0.2218 | 0.1477/ 0.1753 | 0.1505/ 0.1783 | 0.1481/ 0.1886 | 0.1528/ 0.1910 | 0.1479/ 0.1850 | 0.1891/ 0.2236 | 0.1487/ 0.1604 |
| ***No. atoms*** | | | | | | | | | | | | |
| **Protein** | 6359 | 6500 | 6151 | 6657 | 6121 | 6382 | 6412 | 6250 | 6229 | 6175 | 6002 | 6522 |
| **Ligand** | 41 | 46 | 42 | 46 | 42 | 46 | 42 | 46 | 41 | 46 | 40 | 46 |
| **Water** | 184 | 389 | 244 | 437 | 157 | 354 | 330 | 377 | 376 | 343 | 172 | 744 |
| ***Averaged B-factors ($Å^2$)*** | | | | | | | | | | | | |
| **Protein** | 47.22 | 36.58 | 40.90 | 34.25 | 44.08 | 33.71 | 39.07 | 36.83 | 38.93 | 34.33 | 48.41 | 22.73 |
| **Ligand** | 38.82 | 26.19 | 47.31 | 24.68 | 43.05 | 22.71 | 35.15 | 27.55 | 33.19 | 21.09 | 47.62 | 18.10 |
| **Water** | 44.24 | 43.34 | 41.00 | 44.32 | 38.45 | 42.26 | 45.19 | 43.19 | 46.35 | 38.95 | 42.04 | 32.01 |
| ***RMSD*** | | | | | | | | | | | | |
| **bond lengths (Å)** | 0.002 | 0.005 | 0.002 | 0.006 | 0.003 | 0.005 | 0.007 | 0.009 | 0.005 | 0.005 | 0.002 | 0.012 |
| **bond angles (°)** | 0.510 | 0.812 | 0.513 | 0.898 | 0.500 | 0.781 | 0.839 | 0.899 | 0.793 | 0.839 | 0.535 | 1.224 |
| ***Molprobity statistics*** | | | | | | | | | | | | |
| **Ramachand. outliers (%)** | 0.00 | 0.00 | 0.00 | 0.00 | 0.00 | 0.00 | 0.00 | 0.00 | 0.00 | 0.00 | 0.00 | 0.00 |
| **Ramachand. allowed (%)** | 3.40 | 2.39 | 2.90 | 2.64 | 3.03 | 1.90 | 2.39 | 2.14 | 2.27 | 2.02 | 3.38 | 2.90 |
| **Ramachand. favored (%)** | 96.60 | 97.61 | 97.10 | 97.36 | 96.97 | 98.10 | 97.61 | 97.86 | 97.73 | 97.98 | 96.62 | 97.10 |
| **Rotamer outliers (%)** | 0.61 | 0.74 | 1.11 | 1.72 | 0.64 | 1.36 | 1.36 | 1.25 | 1.25 | 0.94 | 0.33 | 0.15 |
| **MolProbity clashscore** | 0.72 | 1.56 | 1.15 | 1.43 | 1.24 | 1.03 | 1.81 | 1.46 | 2.03 | 1.23 | 1.60 | 3.16 |

[a] Highest resolution shell is shown in parentheses.

**Supplementary Table 7**. Hinge movement analysis for AAT variants crystallized here

| DynDom Results | WT (8E9P *vs.* 8E9K) [a] | VFCS (8E9R *vs.* 8E9S) [a] | AIFS (8E9C *vs.* 8E9D) [a] |
|---|---|---|---|
| **Fixed Domain Residues** | 36–320, 340–342, 345–348 | 36–318, 340–348 | 30–318, 339–348 |
| **Moving Domain Residues** | 21–32, 321–339, 343–344, 349–394 | 25–35, 319–339, 349–394 | 29, 319–338, 349–394 |
| **Unassigned Residues** | 1–20, 33–35, 395–396 | 1–24, 395–396 | 1–28, 395–396 |
| **Angle of rotation (°)** | 4.6 | 2.6 | 5.9 |
| **Translation along axis (Å)** | –0.3 | –0.2 | –0.3 |
| **Closure (%)** | 92.5 | 96.0 | 92.8 |
| **Bending Residues** [b] | 30–38, 320–321, 338–349 | 35–36, 318–319, 339–341, 343–352 | 29–46, 318–319, 338–341, 348–352 |

[a] For WT, VFCS, and AIFS, domain movement between the open and closed states was observed for chain A. Hinge movement was not detected for HEX, VFIT, or VFIY. All analyses performed using DynDom (*35*).

**Supplementary Table 8.** Conformational equilibrium constants of AAT variants at various temperatures

| | $K_{eq}$ [a] | | | | | | |
|---|---|---|---|---|---|---|---|
| **Enzyme** | **278 K** | **283 K** | **288 K** | **283 K** | **298 K** | **303 K** | **308 K** |
| **HEX** | 24.093 | 9.047 | 4.301 | 3.139 | 2.190 | 1.730 | 1.482 |
| **VFIT** | 3.002 | 1.380 | 0.790 | 0.494 | 0.368 | 0.279 | 0.231 |
| **VFIY** | 2.058 | 0.810 | 0.425 | 0.219 | 0.124 | 0.066 | 0.046 |
| **AIFS** | 0.157 | 0.521 | 0.795 | 1.570 | 2.308 | 2.715 | 12.735 |

[a] Equilibrium constants were calculated from the ratio of peaks observed by $^{19}F$ NMR at various temperatures (Supplementary Figure 13). For HEX, VFIT, and VFIY, equilibrium constants are reported for the closed/open conformational transition ($K_{eq}$ = closed/open). For AIFS, equilibrium constants are reported for conformational states corresponding to alternate open conformations.

**Supplementary Table 9.** Crystallographic data and refinement statistics for structures at 303 K

| | **WT** | **HEX** | **VFIT** |
|---|---|---|---|
| **Maleate** | (–) | (–) | (–) |
| **PDB ID** | 8E9T | 8E9U | 8E9V |
| **Data collection** [a] | | | |
| **Temp. (K)** | 303 | 303 | 303 |
| **Resolution (Å)** | 124.73–2.13 | 124.43–1.94 | 124.71–2.01 |
| **Space group** | P 63 | P 63 | P 63 |
| ***Cell params.*** | | | |
| **a b c (Å)** | 144.03 144.03 81.26 | 143.66 143.66 81.54 | 144.01 144.01 81.58 |
| **α β γ (°)** | 90 90 120 | 90 90 120 | 90 90 120 |
| **Chains per asymm. unit** | 2 | 2 | 2 |
| **$R_{pim}$** | 0.091 (0.993) | 0.093 (1.860) | 0.083 (0.823) |
| **$CC_{1/2}$** | 0.993 (0.263) | 0.997 (0.334) | 0.993 (0.357) |
| **I/σI** | 6.0 (0.5) | 7.0 (0.6) | 5.5 (0.5) |
| **Complete. (%)** | 100.0 (100.0) | 99.3 (98.7) | 100.0 (100.0) |
| **Multiplicity** | 5.1 (4.8) | 10.2 (10.4) | 5.0 (4.9) |
| **Wilson B-factor (Å$^2$)** | 30.940 | 29.110 | 27.850 |
| **# unique reflections** | 53790 (2679) | 70417 (3433) | 64155 (3220) |
| **Refinement** | | | |
| **R work/free** | 0.1715/ 0.1997 | 0.1473/ 0.1776 | 0.1684/ 0.1961 |
| ***No. atoms*** | | | |
| **Protein** | 6186 | 6275 | 6197 |
| **Ligand** | 41 | 42 | 42 |
| **Water** | 175 | 265 | 223 |
| ***Averaged B-factors (Å$^2$)*** | | | |
| **Protein** | 47.39 | 43.29 | 41.92 |
| **Ligand** | 39.50 | 43.88 | 35.10 |
| **Water** | 45.51 | 47.10 | 44.01 |
| ***RMSD*** | | | |
| **bond lengths (Å)** | 0.002 | 0.007 | 0.002 |
| **bond angles (°)** | 0.504 | 1.008 | 0.517 |
| ***Molprobity statistics*** | | | |
| **Ramachand. outliers (%)** | 0.00 | 0.00 | 0.00 |
| **Ramachand. allowed (%)** | 2.28 | 2.40 | 2.41 |
| **Ramachan. favored (%)** | 97.72 | 97.60 | 97.59 |
| **Rotamer outliers (%)** | 1.26 | 1.70 | 0.90 |
| **MolProbity clashscore** | 1.15 | 1.85 | 1.57 |

[a] Highest resolution shell is shown in parentheses.

**Supplementary Table 10.** Amino-acid sequences of AAT variants

| Enzyme | # mutations from WT | # mutations from HEX | Sequence[a] |
|---|---|---|---|
| **WT** | – | 6 | MAHHHHHHVGTFENITAAPADPILGLADLFRADERPGKINLGIGVYKDETGKTPVLTSVK<br>KAEQYLLENETTKNYLGIDGIPEFGRCTQELLFGKGSALINDKRARTAQTPGGTGALRVA<br>ADFLAKNTSVKRVWVSNPSWPNHKSVFNSAGLEVREYAYYDAENHTLDFDALINSLNEAQ<br>AGDVVLFHGCCHNPTGIDPTLEQWQTLAQLSVEKGWLPLFDFAYQGFARGLEEDAEGLRA<br>FAAMHKELIVASSYSKNFGLYNERVGACTLVAADSETVDRAFSQMKAAIRANYSNPPAHG<br>ASVVATILSNDALRAIWEQELTDMRQRIQRMRQLFVNTLQEKGANRDFSFIIKQNGMFSF<br>SGLTKEQVLRLREEFGVYAVASGRVNVAGMTPDNMAPLCEAIVAVL |
| **HEX** | 6 | – | MAHHHHHHVGTFENITAAPADPILGLADLFRADERPGKINLGIG**L**Y**Y**DETGK**I**PVLTSVK<br>KAEQYLLENETTK**L**YLGIDGIPEFGRCTQELLFGKGSALINDKRARTAQTPGG**S**GALRVA<br>ADFLAKNTSVKRVWVSNPSWPNHKSVFNSAGLEVREYAYYDAENHTLDFDALINSLNEAQ<br>AGDVVLFHGCCHNPTGIDPTLEQWQTLAQLSVEKGWLPLFDFAYQGFARGLEEDAEGLRA<br>FAAMHKELIVASSYSKNFGLYNERVGACTLVAADSETVDRAFSQMKAAIRANYS**S**PPAHG<br>ASVVATILSNDALRAIWEQELTDMRQRIQRMRQLFVNTLQEKGANRDFSFIIKQNGMFSF<br>SGLTKEQVLRLREEFGVYAVASGRVNVAGMTPDNMAPLCEAIVAVL |
| **VFIT** | 3 | 5 | MAHHHHHHVGTFENITAAPADPILGLADLFRADERPGKINLGIGVY**F**DETGK**I**PVLTSVK<br>KAEQYLLENETTK**T**YLGIDGIPEFGRCTQELLFGKGSALINDKRARTAQTPGGTGALRVA<br>ADFLAKNTSVKRVWVSNPSWPNHKSVFNSAGLEVREYAYYDAENHTLDFDALINSLNEAQ<br>AGDVVLFHGCCHNPTGIDPTLEQWQTLAQLSVEKGWLPLFDFAYQGFARGLEEDAEGLRA<br>FAAMHKELIVASSYSKNFGLYNERVGACTLVAADSETVDRAFSQMKAAIRANYSNPPAHG<br>ASVVATILSNDALRAIWEQELTDMRQRIQRMRQLFVNTLQEKGANRDFSFIIKQNGMFSF<br>SGLTKEQVLRLREEFGVYAVASGRVNVAGMTPDNMAPLCEAIVAVL |
| **VFIY** | 3 | 5 | MAHHHHHHVGTFENITAAPADPILGLADLFRADERPGKINLGIGVY**F**DETGK**I**PVLTSVK<br>KAEQYLLENETTK**Y**YLGIDGIPEFGRCTQELLFGKGSALINDKRARTAQTPGGTGALRVA<br>ADFLAKNTSVKRVWVSNPSWPNHKSVFNSAGLEVREYAYYDAENHTLDFDALINSLNEAQ<br>AGDVVLFHGCCHNPTGIDPTLEQWQTLAQLSVEKGWLPLFDFAYQGFARGLEEDAEGLRA<br>FAAMHKELIVASSYSKNFGLYNERVGACTLVAADSETVDRAFSQMKAAIRANYSNPPAHG<br>ASVVATILSNDALRAIWEQELTDMRQRIQRMRQLFVNTLQEKGANRDFSFIIKQNGMFSF<br>SGLTKEQVLRLREEFGVYAVASGRVNVAGMTPDNMAPLCEAIVAVL |
| **VFCS** | 3 | 6 | MAHHHHHHVGTFENITAAPADPILGLADLFRADERPGKINLGIGVY**F**DETGK**C**PVLTSVK<br>KAEQYLLENETTK**S**YLGIDGIPEFGRCTQELLFGKGSALINDKRARTAQTPGGTGALRVA<br>ADFLAKNTSVKRVWVSNPSWPNHKSVFNSAGLEVREYAYYDAENHTLDFDALINSLNEAQ<br>AGDVVLFHGCCHNPTGIDPTLEQWQTLAQLSVEKGWLPLFDFAYQGFARGLEEDAEGLRA<br>FAAMHKELIVASSYSKNFGLYNERVGACTLVAADSETVDRAFSQMKAAIRANYSNPPAHG<br>ASVVATILSNDALRAIWEQELTDMRQRIQRMRQLFVNTLQEKGANRDFSFIIKQNGMFSF<br>SGLTKEQVLRLREEFGVYAVASGRVNVAGMTPDNMAPLCEAIVAVL |
| **AIFS** | 4 | 6 | MAHHHHHHVGTFENITAAPADPILGLADLFRADERPGKINLGIG**A**Y**I**DETGK**F**PVLTSVK<br>KAEQYLLENETTK**S**YLGIDGIPEFGRCTQELLFGKGSALINDKRARTAQTPGGTGALRVA<br>ADFLAKNTSVKRVWVSNPSWPNHKSVFNSAGLEVREYAYYDAENHTLDFDALINSLNEAQ<br>AGDVVLFHGCCHNPTGIDPTLEQWQTLAQLSVEKGWLPLFDFAYQGFARGLEEDAEGLRA<br>FAAMHKELIVASSYSKNFGLYNERVGACTLVAADSETVDRAFSQMKAAIRANYSNPPAHG<br>ASVVATILSNDALRAIWEQELTDMRQRIQRMRQLFVNTLQEKGANRDFSFIIKQNGMFSF<br>SGLTKEQVLRLREEFGVYAVASGRVNVAGMTPDNMAPLCEAIVAVL |

[a] Sequence for WT was obtained from Uniprot (P00509). Mutations from wild-type (WT) AAT are highlighted in bold and underlined. All sequences contain a His-tag at the N-terminus.

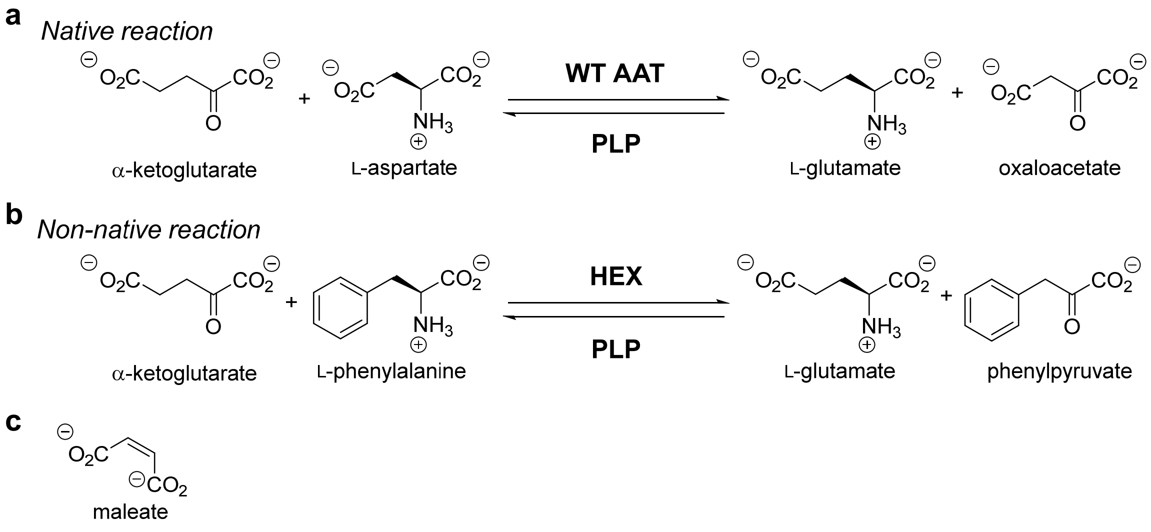


**Supplementary Figure 1. Transamination reactions catalyzed by *E. coli* aspartate aminotransferase (AAT).** (a) Wild-type (WT) AAT natively catalyzes the reversible transamination of dicarboxylate substrates using the pyridoxal 5’-phosphate (PLP) cofactor. (b) The AAT hexamutant (HEX) can also efficiently catalyze transamination of the non-native substrate L-phenylalanine. (c) AAT is inhibited by L-aspartate analogue maleate.

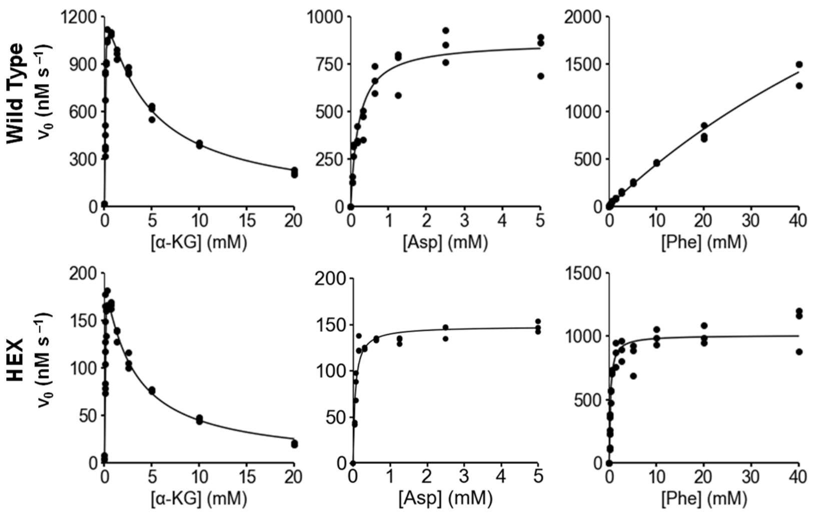


**Supplementary Figure 2. Steady-state kinetics of wild-type AAT and HEX.** Michaelis-Menten plots of initial rates (normalized to enzyme quantity) as a function of substrate concentrations are shown. α-KG, Asp, and Phe indicate α-ketoglutarate, L-aspartate, and L-phenylalanine, respectively. All experiments were performed in triplicate. For α-KG, fitting of the kinetic data was done with a rate equation that takes into account substrate inhibition (Methods).

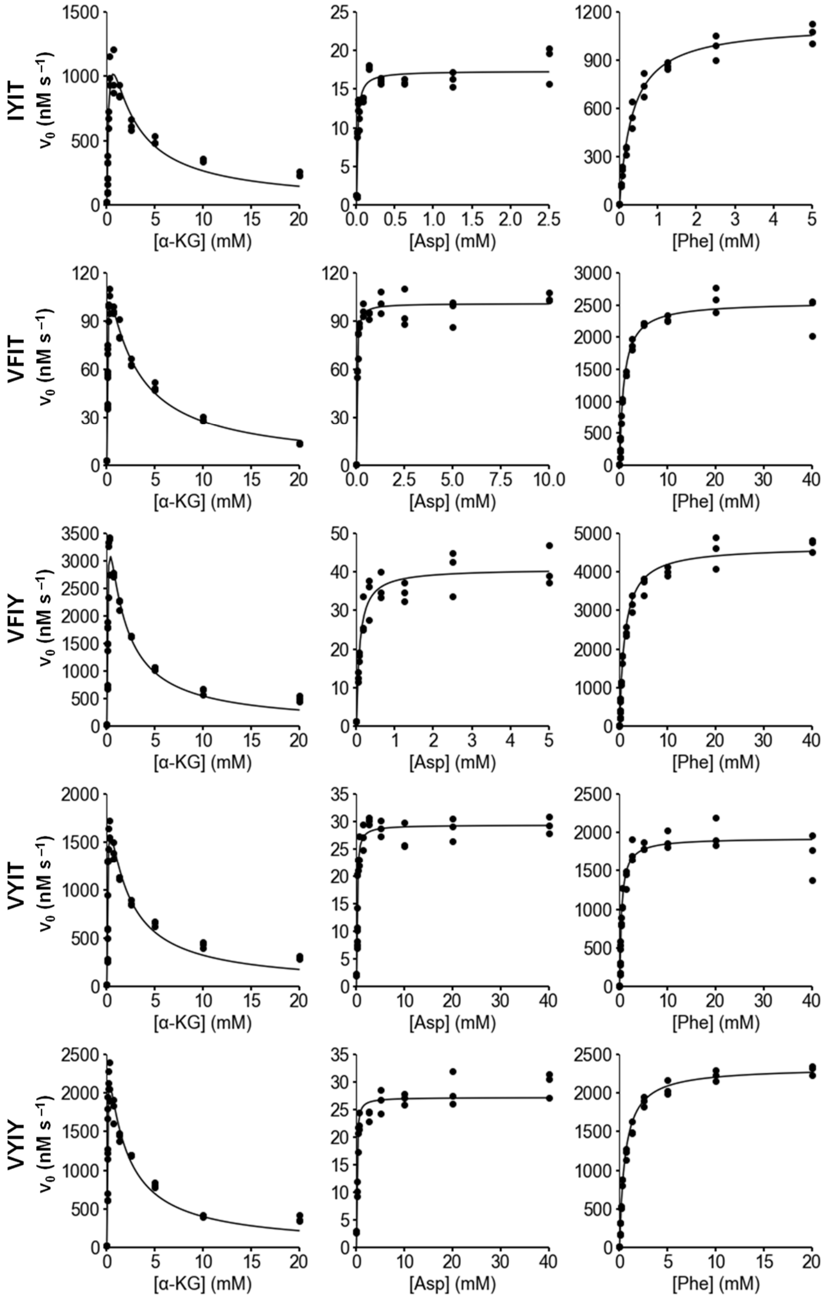


**Supplementary Figure 3. Steady-state kinetics of Closed library mutants.** Michaelis-Menten plots of initial rates (normalized to enzyme quantity) as a function of substrate concentrations are shown. α-KG, Asp, and Phe indicate α-ketoglutarate, L-aspartate, and L-phenylalanine, respectively. All experiments were performed in triplicate. For α-KG, fitting of the kinetic data was done with a rate equation that takes into account substrate inhibition (Methods).

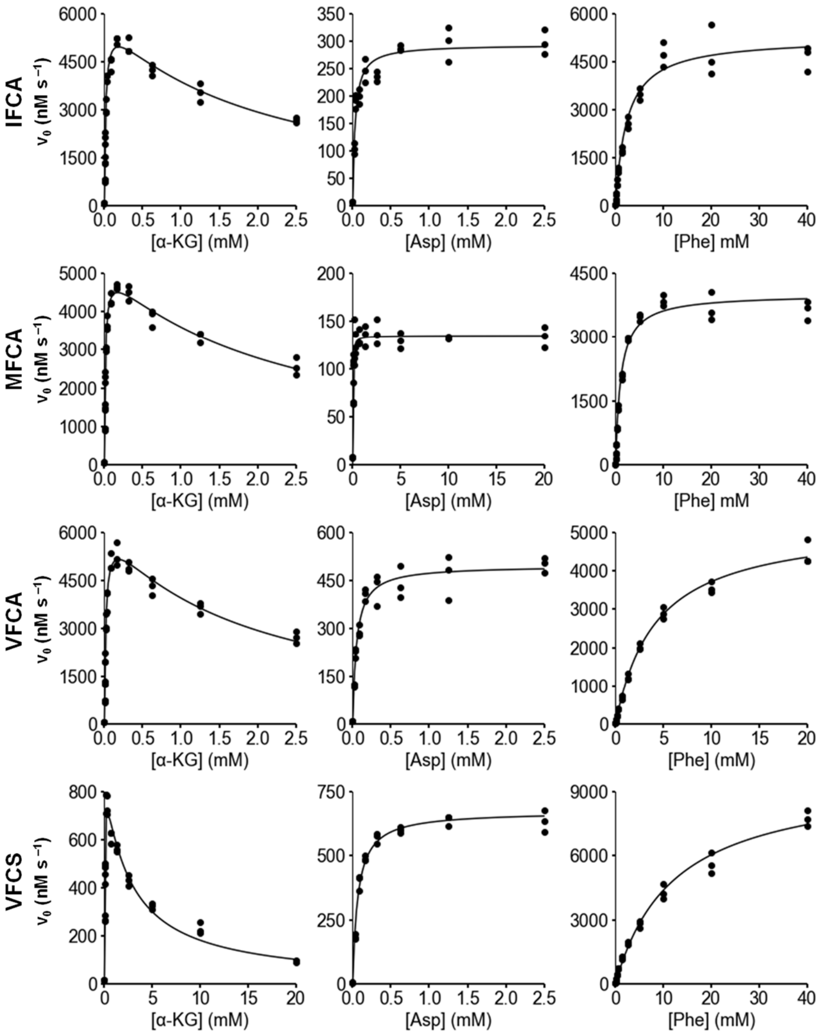


**Supplementary Figure 4. Steady-state kinetics of Open$_{Low}$ library mutants.** Michaelis-Menten plots of initial rates (normalized to enzyme quantity) as a function of substrate concentrations are shown. α-KG, Asp, and Phe indicate α-ketoglutarate, L-aspartate, and L-phenylalanine, respectively. All experiments were performed in triplicate. For α-KG, fitting of the kinetic data was done with a rate equation that takes into account substrate inhibition (Methods).

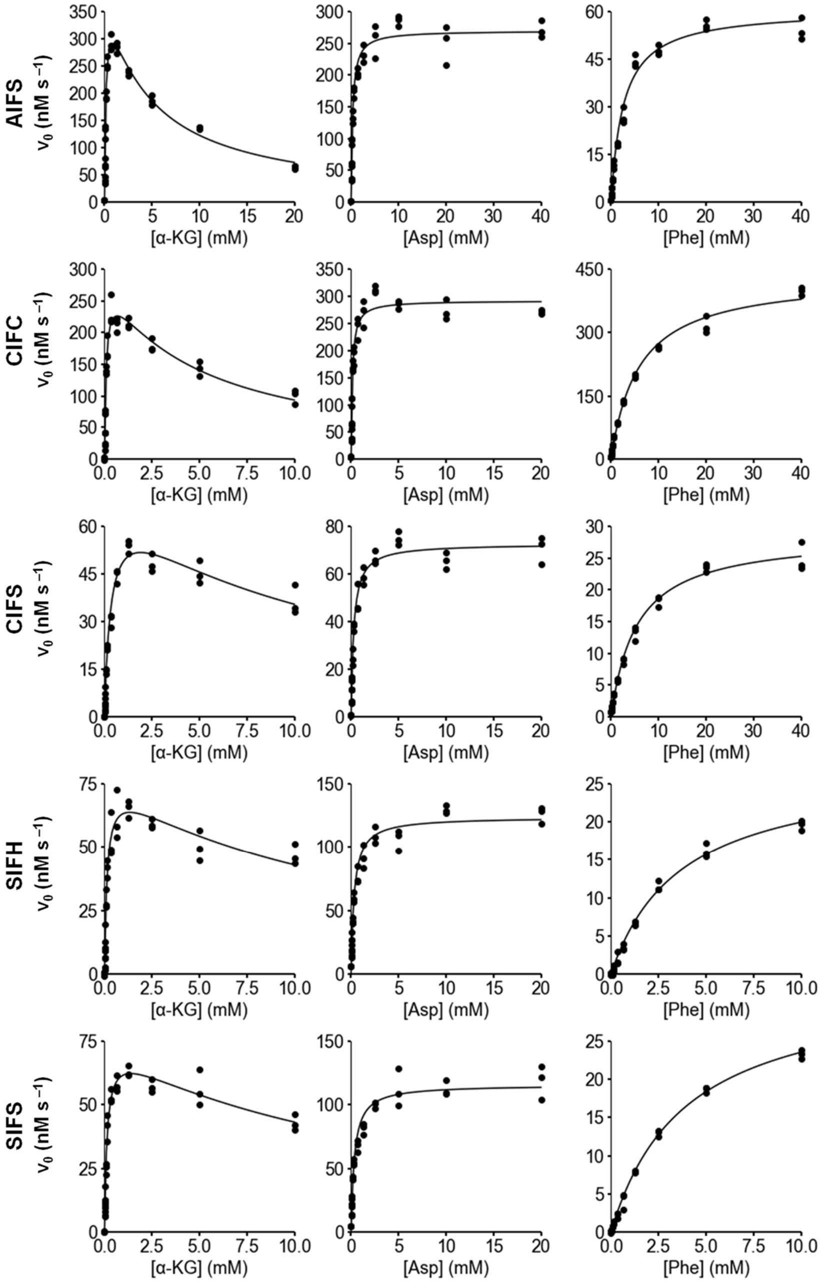


**Supplementary Figure 5. Steady-state kinetics of $Open_{High}$ library mutants.** Michaelis-Menten plots of initial rates (normalized to enzyme quantity) as a function of substrate concentrations are shown. α-KG, Asp, and Phe indicate α-ketoglutarate, L-aspartate, and L-phenylalanine, respectively. All experiments were performed in triplicate. For α-KG, fitting of the kinetic data was done with a rate equation that takes into account substrate inhibition (Methods).

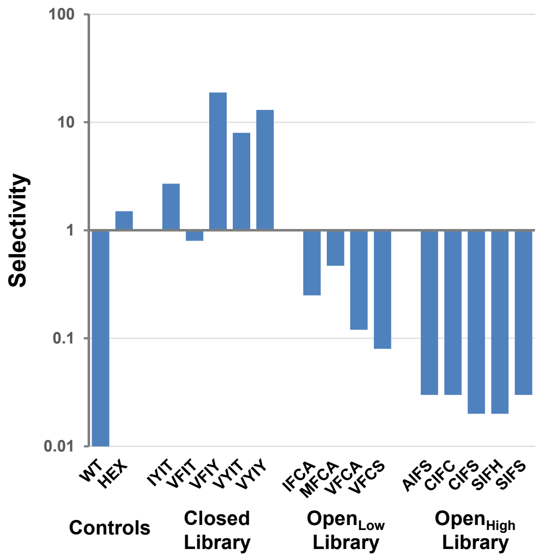


**Supplementary Figure 6. Substrate selectivity of AAT variants.** Selectivity is defined as ($k_{cat}/K_M$ L-phenylalanine) / ($k_{cat}/K_M$ L-aspartate). Although Closed Library mutant VFIT has a selectivity value of 0.8, it is approximately 60-fold more active with the non-native substrate than the wild type (WT).

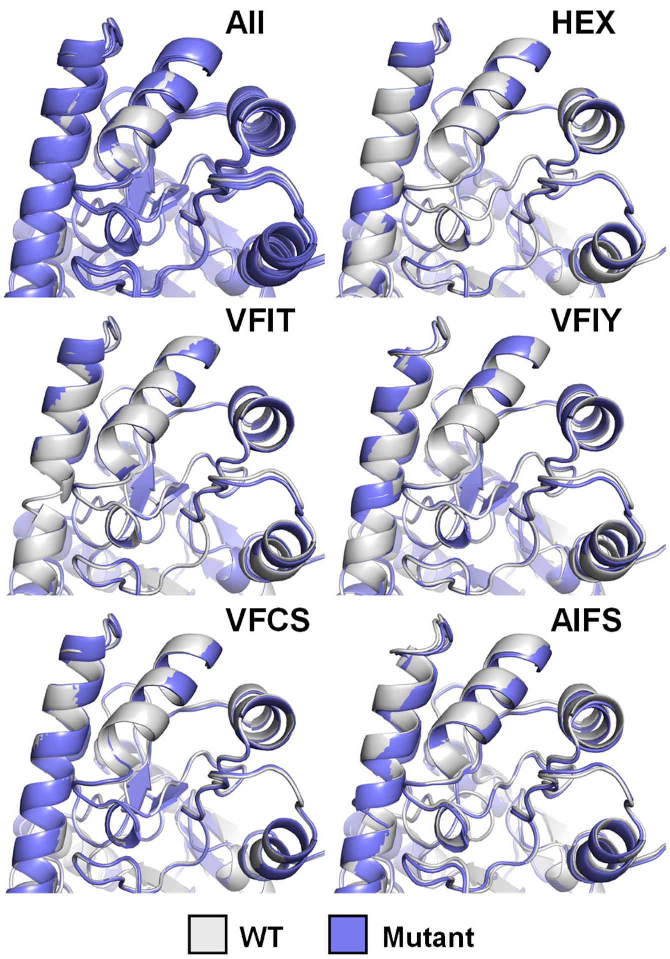


**Supplementary Figure 7. Crystal structures of AAT variants in the maleate-bound form.** Overlay of crystal structures shows that all six variants adopt nearly identical closed conformations in the presence of inhibitor.

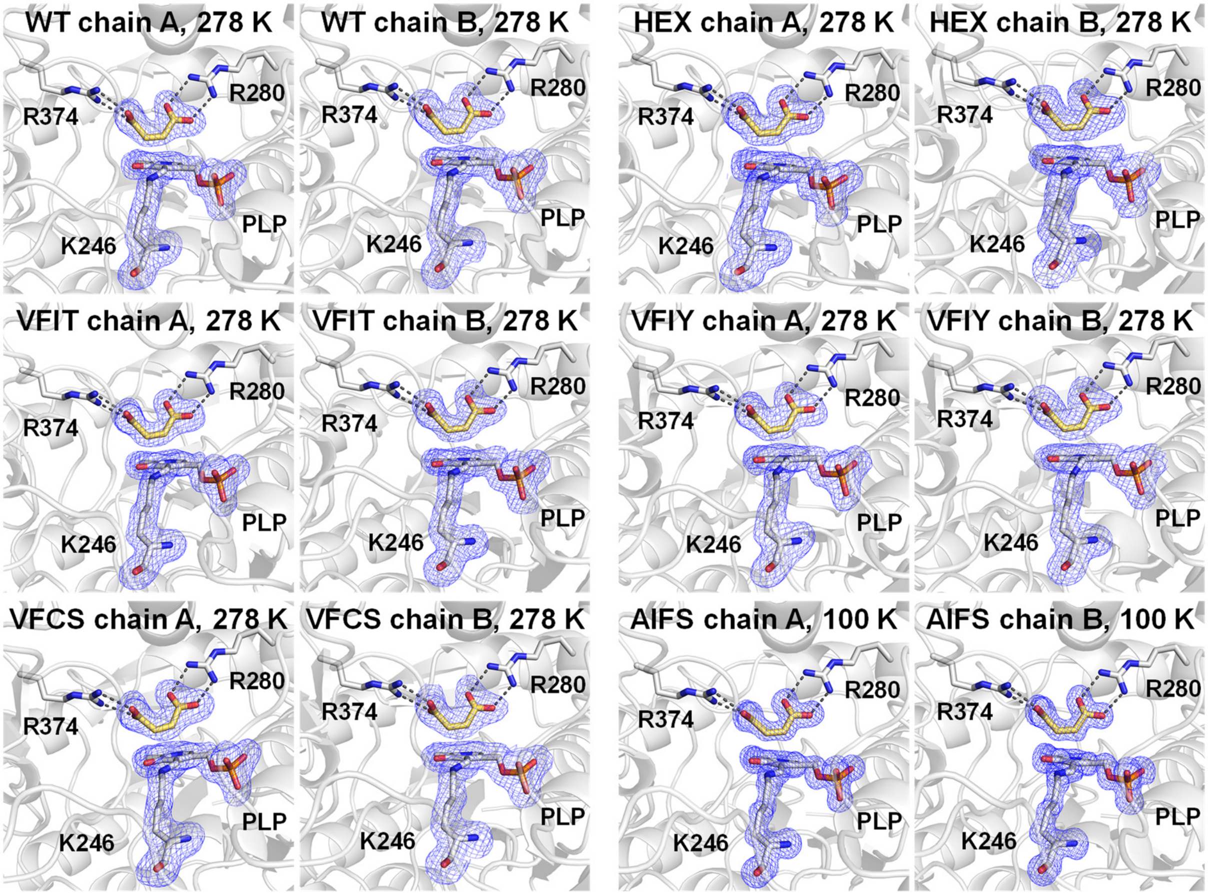


**Supplementary Figure 8. Crystallographic evidence of the presence of bound maleate for all AAT variants.** Presence of bound maleate (yellow), in both chain A and chain B of the enzyme models at various temperatures, is confirmed by an omit map (3.0σ, blue). The electron density supports a binding mode in which maleate forms hydrogen bonds (dashed lines) to both R280 and R374. In all cases, enzymes adopt the internal aldimine form where the catalytic K246 residue forms a Schiff base with the pyridoxal 5'-phosphate (PLP) cofactor.

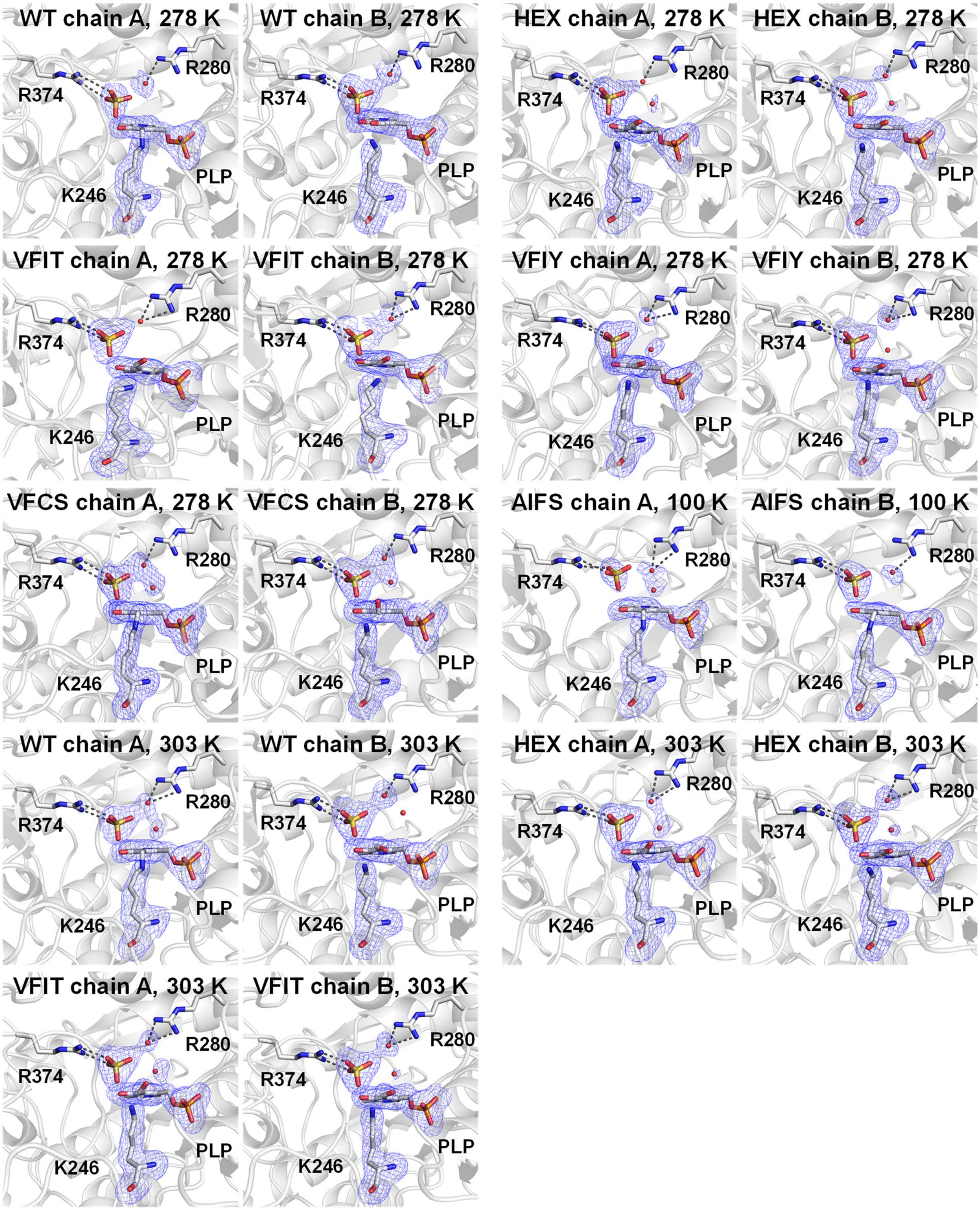


**Supplementary Figure 9. Crystallographic evidence of maleate removal by crystal soaking.** Removal of bound maleate, in both chain A and chain B of the enzyme models at various temperatures, is confirmed by an omit map (3.0σ, blue). The electron density supports replacement of maleate by a sulfate ion from the crystallization buffer and one or more water molecules (red spheres), which form electrostatic interactions or hydrogen bonds (dashed lines) to R280 or R374. The crystal soaking procedure to remove maleate also causes the loss of the covalent bond between the catalytic K246 residue and pyridoxal 5'-phosphate (PLP) cofactor in one or both subunits.

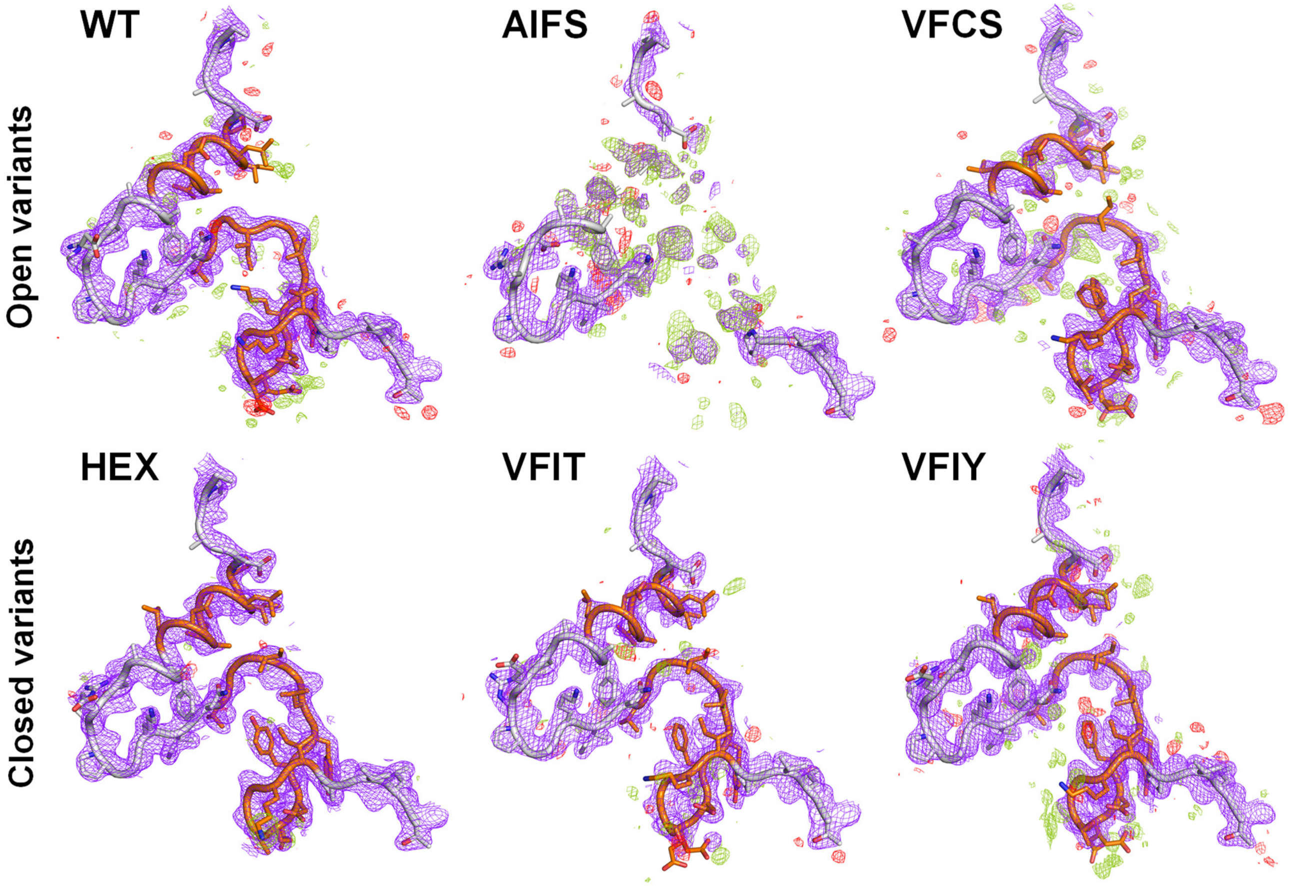


**Supplementary Figure 10. Conformational heterogeneity in $Open_{High}$ library mutant AIFS.** Electronic density of Chain A residues Ala8–Thr47 in all AAT variants in their ligand-free forms is shown, with 2mFo-DFc electron density maps contoured at 1 σ (blue mesh) and mFo-DFc difference density maps contoured at ± 3 σ (green/red mesh). There is much heterogeneity in electron density around residues Pro12–Leu19 and Leu31–Thr43 (orange) in AIFS but not other variants.

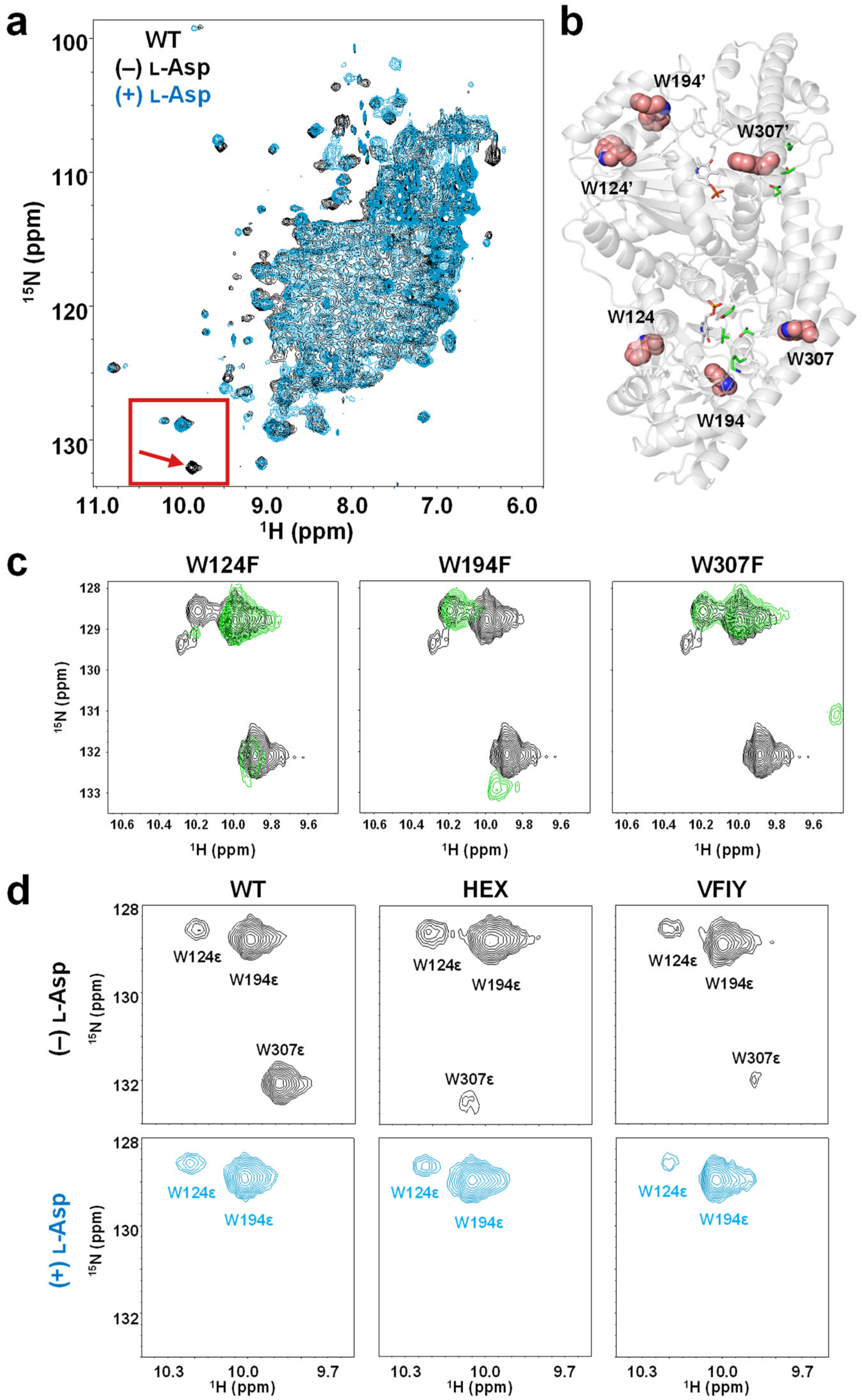


**Supplementary Figure 11. $^{1}$H-$^{15}$N HSQC spectra of selected AAT variants.** (a) Addition of 5 mM donor substrate to wild-type (WT) AAT causes the shifting of peaks throughout the spectrum and disappearance of a peak (indicated by arrow) in the Trp side-chain region (boxed). (b) Trp residues whose side-chain amides yield peaks in the Trp side-chain region are shown as spheres (salmon). The PLP cofactor bound at the active site and designed residues (V35, K37, T43, N64) are shown as white and green sticks, respectively. (c) Overlay of WT spectrum (black) on those of single point mutants (green) allowed assignment of Trp indole NH peaks. All spectra were measured in the absence of substrate. (d) The side-chain peak of W307 disappears when 5 mM L-aspartate is added. In HEX and the closed library mutant VFIY, this peak is already substantially broadened in the absence of substrate, providing initial evidence that the conformational equilibrium in these variants favours the closed state.

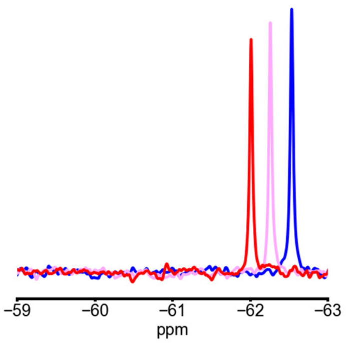


**Supplementary Figure 12. $^{19}$F NMR spectra of 4-trifluoromethyl-L-phenylalanine at various temperatures.** Blue, purple, and red lines correspond to spectra at 278, 293, and 308 K, respectively.

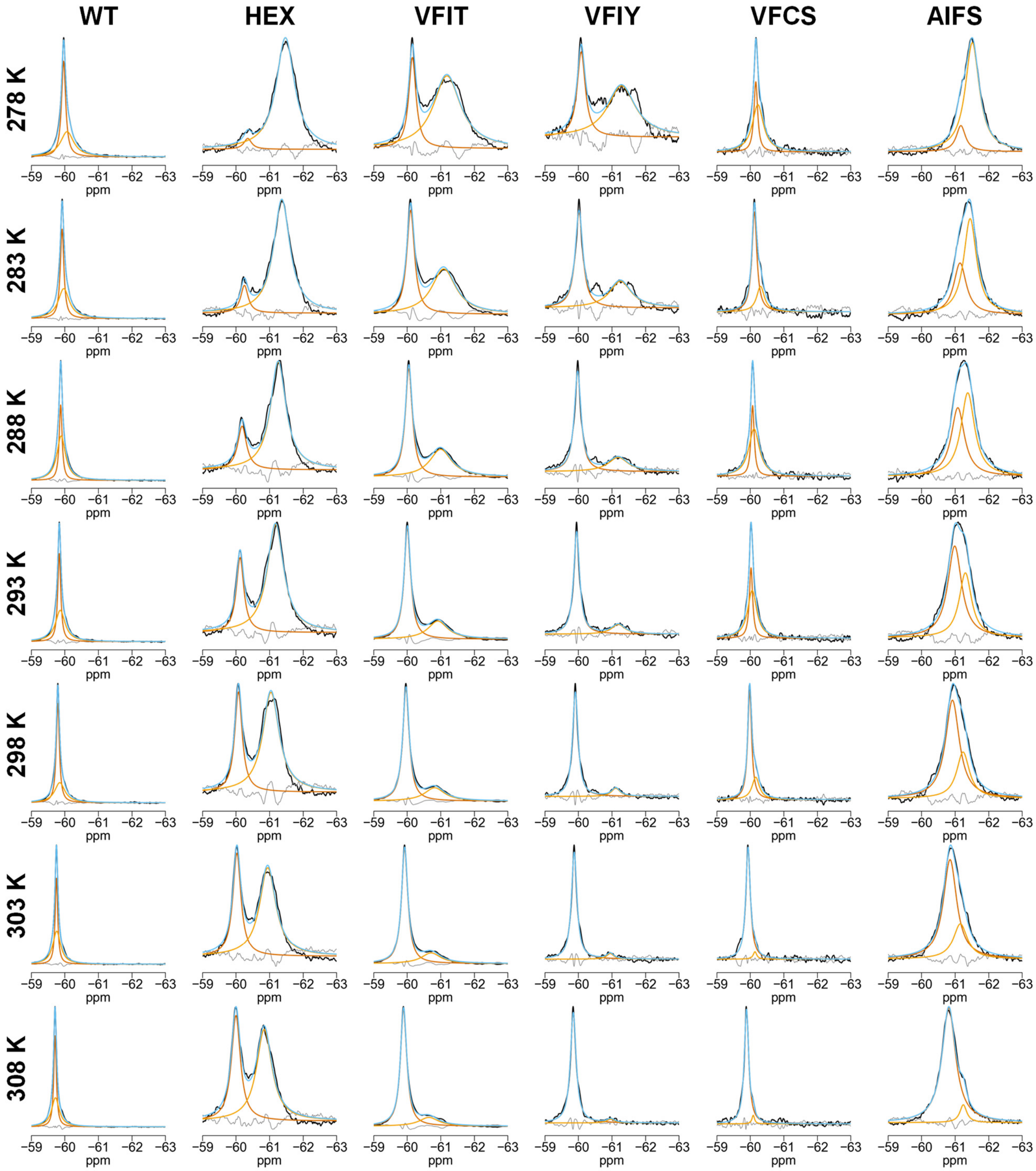


**Supplementary Figure 13. Deconvolution of $^{19}$F NMR spectra at various temperatures.** The Lorentzian function was used to fit two peaks (light and dark orange) to the experimental NMR spectra (black) of various AATs. Fitted spectra and residuals are colored blue and grey, respectively. For WT and VFCS, deconvolution of spectra as two separate peaks did not yield meaningful results.

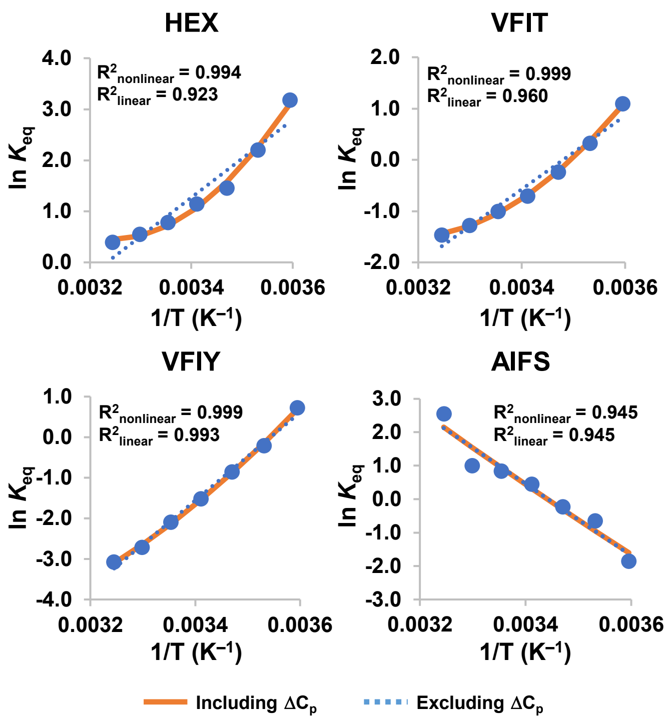


**Supplementary Figure 14. Thermodynamic analysis of conformational exchange.** The van't Hoff equation including (nonlinear) or excluding (linear) changes to heat capacity ($\Delta C_p$) was used to fit equilibrium constants as a function of temperature. Coefficients of determination ($R^2$) close to unity for HEX, VFIT, VFIY, and AIFS confirm that these proteins are undergoing exchange. For HEX, VFIT, and VFIY, nonlinear fitting improves $R^2$ values, which is not the case for AIFS. For conformational exchange of AIFS between alternate open conformations (linear fit), we calculated ΔG, ΔH, and ΔS values (278 K) of 0.83 kcal mol$^{-1}$, 21.4 kcal mol$^{-1}$, and 0.074 kcal mol$^{-1}$ K$^{-1}$, respectively.

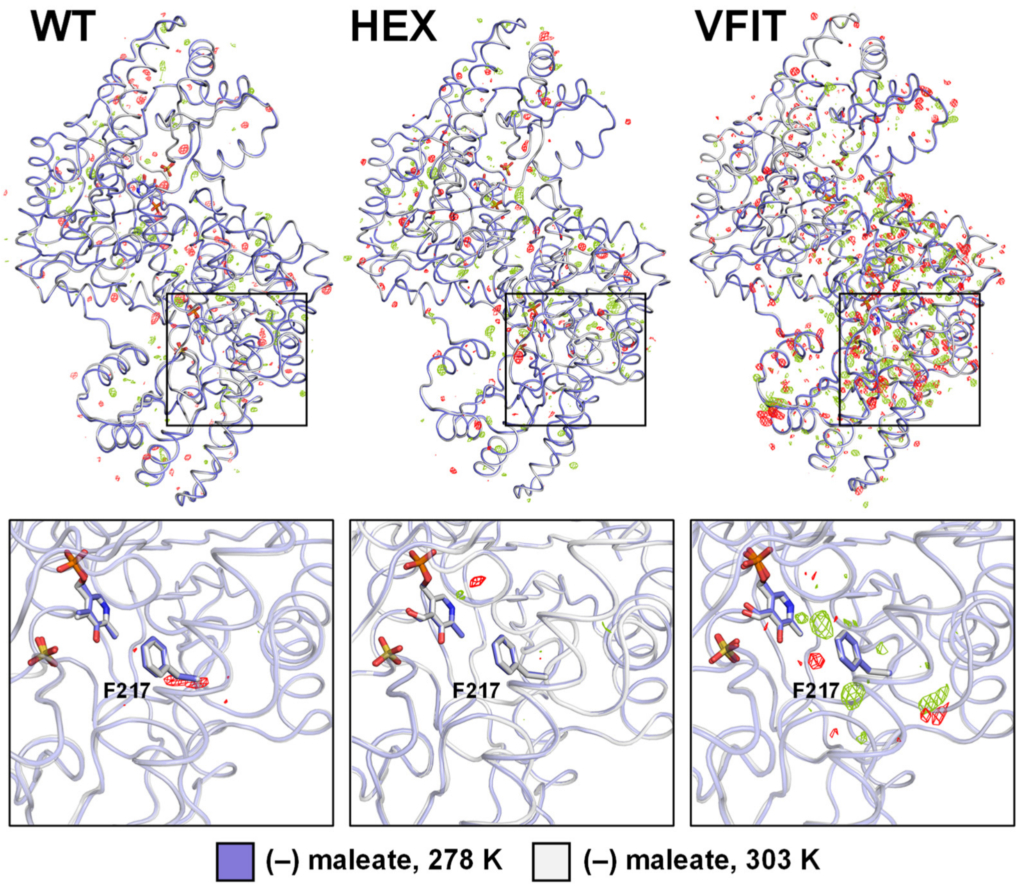


**Supplementary Figure 15. Fo-Fo difference maps reveal increased local conformational changes in VFIT when temperature of crystal is increased from 278 K to 303 K.** Top: Isomorphous Fo-Fo difference electron density map contoured at ± 3 σ (green/red mesh) for the 303 K dataset (white) minus the 278 K dataset (blue) in the absence of maleate. Bottom: Difference density (± 3 σ) within a 5 Å radius of the F217 residue (sticks) where the 4-trifluoromethyl-L-phenylalanine amino acid was incorporated to enable multitemperature $^{19}$F NMR spectroscopy. There is more difference density throughout chain A in the VFIT data set, even though the backbone conformation in crystal does not change substantially upon heating. The pyridoxal 5'-phosphate cofactor and sulfate ion that occupies the active site in the absence of maleate are shown as sticks. In all cases, isomorphous difference maps were generated using high- and low-resolution cut-offs of 2.31 and 20.0 Å, respectively.

### Materials and Methods

*Structure preparation and ensemble generation.* Crystal structures of wild-type *Escherichia coli* AAT in its internal aldimine form (PDB ID: 1ARS (*12*)) or complexed with *N*-phosphopyridoxyl-L-glutamic acid (PDB ID: 1X28 (*15*)) were used to model the open and closed states, respectively. To eliminate biases that could arise during ensemble generation from the presence of substrate bound in the active site, we deleted coordinates for the *N*-phosphopyridoxyl-L-glutamic acid and catalytic K246 residue in the 1X28 structure and replaced them with coordinates for K246 and the pyridoxal 5'-phosphate (PLP) cofactor extracted from the 1ARS structure. These structures were then prepared for ensemble generation using the Molecular Operating Environment (MOE) software (*36*). Hydrogens were added with the Protonate3D utility and manually adjusted to ensure that the protonation states of PLP and K246 were consistent with the aminotransferase catalytic mechanism (*37*). The resulting structures were then solvated in a rectangular box of water with counter ions ($Na^+$ and $Cl^-$) under periodic boundary conditions with a box cut-off of 6 Å, and energy-minimized by conjugate gradient energy minimization to a root mean square gradient < 7 kcal $mol^{-1}$ $Å^{-1}$ using the AMBER99 force field (*38*) with a combined explicit solvent and implicit reaction field solvent model set up using the MOE software package. These structures were used as input templates to generate backbone ensembles with the PertMin algorithm (*17, 18*). Briefly, two 50-member PertMin ensembles were created by randomly perturbing the coordinates of all heavy atoms of the two prepared AAT structures by ± 0.001 Å along each Cartesian coordinate axis, and energy minimizing them using a truncated Newton (*39*) minimization algorithm for 100 iterations. The "Open" and "Closed" ensembles thus obtained displayed diversities (i.e. average backbone root mean square deviations between pairs of ensemble members) of 0.29 ± 0.03 Å and 0.32 ± 0.03 Å, respectively, and backbone root mean square deviations from the starting structure of 0.51 ± 0.02 Å.

*Computational protein design.* All calculations were performed using the Phoenix protein design software (*20, 40*) with the fast and accurate side-chain topology and energy refinement (FASTER) algorithm (*41*) for sequence optimization. The 2002 backbone-dependent Dunbrack rotamer library (*42*) with expansions of ± 1 standard deviation around $\chi_1$ and $\chi_2$ was used to provide side-chain conformations of AAT residues to be threaded onto each fixed backbone template. Sequences were scored using the Phoenix energy function, a five-term potential energy function consisting of a Lennard-Jones 12–6 van der Waals term from the Dreiding II force field (*43*) with atomic radii scaled by 0.9, a direction-dependent hydrogen bond term with a well depth of 8.0 kcal mol$^{-1}$ and an equilibrium donor-acceptor distance of 2.8 Å (*44*), an electrostatic energy term modelled using Coulomb's law with a distance-dependent dielectric of 10, an occlusion-based solvation penalty term (*20*), and a secondary structural propensity term (*45*). Sidechain rotamers of residues 35, 37, 43, and 64 were optimized on each backbone template using all proteinogenic amino acids with the exception of proline. Sidechain rotamers of residues within 5 Å of the designed residues were also optimized but their identities were not changed. The searched sequence space thus consisted of 130,321 ($19^4$) sequences, resulting in >13 million individual sequence energies (130,321 sequences × 50 backbones × 2 ensembles).

Boltzmann weighted average potential energies at 300 K were computed for each sequence on each ensemble, yielding energy values for the Closed and Open states ($E_{closed}$ and $E_{open}$). Energy differences between conformational states ($\Delta E = E_{closed} - E_{open}$) were then computed for use in library design. To avoid favoring sequences displaying unfavorable $E_{closed}$ and/or $E_{open}$ values resulting in favorable ΔE (e.g., high, low, or high absolute ΔE values for Closed, $Open_{Low}$, or $Open_{High}$ libraries, respectively), which would be expected to be unstable, sequences whose $E_{closed}$ and/or $E_{open}$ value fell outside of the 75$^{th}$ percentile were discarded.

*Library design.* Library design was performed with the CLEARSS algorithm (*19*) using as input the ΔE values obtained as described above. For a specific library size configuration, which is the specific number of amino acids at each position in the protein (e.g., 4 amino acids at position 1, 3 amino acids at position 2, etc.), the highest probability set of amino acids at each position were included in the library. To identify the optimal library size configuration, all configurations that lead to a combinatorial library of target sizes (in this case, 20 ± 4 sequences) were scored by taking the sum of all partition functions of the chosen amino acid sets over all positions, and the highest scoring library was selected. $E_{closed}$, $E_{open}$, and ΔE values for mutants from each library are reported on Supplementary Table 2.

*Chemicals.* All reagents used were of the highest available purity. Synthetic oligonucleotides were purchased from Eurofins MWG Operon. Restriction enzymes and DNA-modifying enzymes were purchased from New England Biolabs. Ni-NTA agarose resin was purchased from Bio-Rad Laboratories. All aqueous solutions were prepared using water purified with a Barnstead Nanopure Diamond system.

*Mutagenesis.* The wild-type *E. coli* AAT gene (Uniprot ID: P00509) with an N-terminal His-tag cloned into plasmid pET-45b (Novagen) via the *Nco*I/*Pac*I restriction sites (*46*) was a generous gift from Michael D. Toney (University of California, Davis). Mutations were introduced into the AAT gene by overlap extension mutagenesis (*47*) using VentR DNA Polymerase. Briefly, external primers containing *Nde*I or *Bam*HI restriction sites were used in combination with sets of complementary pairs of oligonucleotides containing mutated codons (individual codons for HEX and single point mutants, codon mixtures for the Closed, $Open_{Low}$, and $Open_{High}$ libraries) in individual polymerase chain reactions (PCRs). The resulting overlapping fragments were gel-purified (Omega Biotek) and recombined by overlap extension PCR. The resulting amplicons were digested with *Nde*I/*Bam*HI, gel-purified, and ligated into the pET-11a expression vector (Novagen) with T4 DNA ligase. pBAD vectors (Invitrogen) harbouring selected AAT genes flanked by *Nco*I/*Eco*RI restriction sites were prepared using a similar procedure. All constructs were verified by sequencing the entire open reading frame. Amino-acid sequences of all AAT variants are listed in Supplementary Table 10.

*Preparation of clarified cell lysates.* DNA libraries prepared as described above were transformed into chemically competent *E. coli* BL21-Gold (DE3) cells (Agilent). Colonies (180 per library) were picked into individual wells of V96 MicroWell polypropylene plates (Nunc) containing 300 µL of lysogeny broth (LB) supplemented with 100 µg $mL^{-1}$ ampicillin and 10% glycerol. The plates were covered with a sterile breathable rayon membrane (VWR) and incubated overnight at 37 °C with shaking. After incubation, these mother plates were used to inoculate sterile Nunc V96 MicroWell polypropylene plates (''daughter'' plates) containing 300 µL per well of Overnight Express Instant TB medium (Novagen) supplemented with ampicillin. Daughter plates were sealed with breathable membranes and incubated overnight (37 °C, 250 rpm). After incubation, cells were harvested by centrifugation (3000×*g*, 30 min, 4 °C) and pellets were washed twice with phosphate-buffered saline (pH 7.4). Washed cell pellets were resuspended in lysis buffer (100 mM potassium phosphate buffer pH 8.0 containing 1× Bug Buster Protein Extraction Reagent [Novagen], 5 U $mL^{-1}$ Benzonase Nuclease [EMD], and 1 mg $mL^{-1}$ lysozyme). Clarified lysates were collected following centrifugation and stored at 4 °C until used in the screening assay.

*Library screening.* All assays were performed in 200-μL reactions at 37 °C in 100 mM potassium phosphate buffer (pH 8.0). The standard reaction mixture contained final concentrations of either 3 or 40 mM L-phenylalanine, 16 μM PLP, 0.2 mM α-ketoglutarate, 1 U of glutamate dehydrogenase (GDH) from bovine liver (Sigma), and 5 mM $NAD^+$. Plates containing the standard reaction mixture were incubated at 37 °C for 5 min prior to initiation of the reaction by addition of 10 μL of clarified cell lysates prepared as described above. Enzyme reactions were monitored by measuring absorbance of NADH at 340 nm every 12 sec for 30 or 60 min in individual wells of 96-well plates (Greiner Bio-One) using a SpectraMax 384 Plus plate reader (Molecular Devices). The four or five most active variants from each library were selected for further characterization.

*Protein expression and purification.* Selected mutants were expressed and purified as described by Mironov *et al.* (*48*). Briefly, *E. coli* BL21-Gold (DE3) cells harbouring expression vectors containing aminotransferase genes were grown at 37 °C in 500 mL LB medium supplemented with 100 μg $mL^{-1}$ ampicillin until they reached an OD600 of 0.6. Isopropyl β-D-1-thiogalactopyranoside (1 mM) was added to the flasks to induce protein expression, followed by shaking overnight at 16 °C. Cells were harvested by centrifugation, resuspended in 10 mL lysis buffer (5 mM imidazole in 100 mM potassium phosphate buffer, pH 8.0), and lysed with an EmulsiFlex-B15 cell disruptor (Avestin). Proteins were purified by immobilized metal affinity chromatography using Ni–NTA agarose pre-equilibrated with lysis buffer in individual Econo-Pac gravity-flow columns (Bio-Rad). Columns were washed twice, first with 10 mM imidazole in 100 mM potassium phosphate buffer (pH 8.0), and then with the same buffer containing 20 mM imidazole. Bound proteins were eluted with 250 mM imidazole in 100 mM potassium phosphate buffer (pH 8.0) and exchanged into 100 mM sodium phosphate buffer (pH 8.0) using Econo-Pac 10DG desalting pre-packed gravity flow columns (Bio-Rad). For crystallography, proteins were further purified by gel filtration in 20 mM potassium phosphate buffer (pH 7.5) using an

ENrich SEC 650 size-exclusion chromatography column (Bio-Rad). Purified samples were concentrated using Microsep Advance 10K centrifugal devices (Pall) to a final concentration of 170 μM. Protein concentrations were quantified using a modified version of the Bradford assay, where the calibration curve is constructed as a plot of the ratio of the absorbance measurements at 590 and 450 nm versus concentration (*49*).

*Steady-state kinetics.* To measure steady-state kinetics for the α-ketoglutarate acceptor substrate, assays were performed by varying the α-ketoglutarate concentration from 0.002 to 20 mM in the presence of 10–20 mM L-phenylalanine or L-aspartate (Supplementary Table 4), 5 mM $NAD^+$, 16 μM PLP, 1 U GDH, and approximately 10 mU of aminotransferase in 100 mM potassium phosphate buffer (pH 8, 37 °C). To measure steady-state kinetics for the L-aspartate and L-phenylalanine donor substrates, assays were performed by varying the amino-acid concentration from 0.002 to 40 mM in the presence of 0.1563–1.25 mM α-ketoglutarate (Supplementary Table 3), 5 mM $NAD^+$, 16 μM PLP, 1 U GDH, and approximately 10 mU of aminotransferase in 100 mM potassium phosphate buffer (pH 8, 37 °C). The pH of all reaction mixtures was adjusted to 8.0 prior to initiation of the reaction. Enzyme reactions were monitored by measuring absorbance of NADH at 340 nm ($\varepsilon$ = 6220 $M^{-1}$ $cm^{-1}$) every 12 sec for 30 or 60 min in individual wells of 96-well plates (Greiner Bio-One) using a SpectraMax 384 Plus plate reader (Molecular Devices). Path lengths for each well were calculated ratiometrically using the difference in absorbance of potassium phosphate buffer at 900 and 998 nm. Linear phases of kinetic traces were used to measure initial reaction rates. Initial reaction rates at different substrate concentrations were fit to the Michaelis-Menten equation using Python ver.2.7.15 with the *scipy.optimize.curve_fit* function (scipy ver.1.1.0). For mutant and substrate combinations resulting in substrate inhibition, fitting of the kinetic data was done with a rate equation that takes into account this type of inhibition: $v_0 = (v_{max}[S])/(K_M + [S] + [S]^2/K_i)$.

*Preparation of $^{15}N$ labeled proteins.* Proteins for NMR spectroscopy were expressed using M9 minimal expression medium supplemented with 1 g $L^{-1}$ $^{15}N$-labeled ammonium chloride ($^{15}NH_4Cl$) for isotopic enrichment. Cultures were grown at 37 °C with shaking to an optical density at 600 nm of approximately 0.6, after which protein expression was initiated with 1 mM isopropyl β-D-1-thiogalactopyranoside. Following overnight incubation at 16 °C with shaking (275 rpm), cells were harvested by centrifugation and lysed with an EmulsiFlex-B15 cell disruptor (Avestin). Proteins were purified by immobilized metal affinity chromatography as described above, which was followed by gel filtration in 10 mM sodium phosphate buffer (pH 6.0) using an ENrich SEC 650 size exclusion chromatography column (Bio-Rad). Purified samples were concentrated using Amicon Ultracel-10K centrifugal filter units (EMD Millipore).

*Preparation of $^{19}F$ site-specific labeled proteins.* Site-specific incorporation of 4-trifluoromethyl-L-phenylalanine into selected AAT variants was prepared with a protocol adapted from Hammill *et al.* (*25*). Briefly, chemically-competent *E. coli* DH10B cells (Thermo Fisher) harbouring the pDule-4-tfmF A65V S158A plasmid (Addgene plasmid #85484) (*50*), which encodes an orthogonal aminoacyl-tRNA synthetase and cognate amber suppressing tRNA for site-specific incorporation of 4-trifluoromethyl-L-phenylalanine, were transformed with pBad vectors (Invitrogen) containing mutated genes of AAT variants in which an amber stop codon was introduced at residue position 217 to allow direct incorporation of the fluorinated amino acid. Labelled AAT variants were expressed in 2 L of LB containing 100 μg $mL^{-1}$ ampicillin and 15 μg $mL^{-1}$ tetracycline at 37 °C. After cells were grown for 1 h, 468 mg of 4-trifluoromethyl-L-phenylalanine (SynQuest Laboratories) was added to the flask to give a final concentration of 1 mM. Once the cell culture reached an OD600 of 0.6, L-arabinose was added to a final concentration of 0.2% to induce protein expression. Following overnight incubation at 16 °C with shaking, cells were harvested by centrifugation, resuspended in 10 mL lysis buffer, and lysed with an EmulsiFlex-B15 cell disruptor (Avestin). Proteins were then extracted and purified by immobilized metal affinity chromatography, as described above. Elution fractions containing the aminotransferases were concentrated through centrifugation (Pall Microsep Advance Centrifugal Device 10K), resuspended in 10 mM potassium phosphate buffer pH 6.0 and further purified through size exclusion chromatography, as described above. Elution fractions containing the purified aminotransferase were combined and concentrated through centrifugation.

*$^{1}H$-$^{15}N$ heteronuclear single quantum coherence spectroscopy.* $^{15}$N-labeled AAT samples for NMR (wild type, HEX, VFIY, and single point mutants Trp124Phe, Trp194Phe, Trp307Phe) consisted of 0.2–0.6 mM protein in 10 mM sodium phosphate buffer (pH 7.4), 10 μM EDTA, 0.02% sodium azide, and 10% $D_2O$. HSQC experiments were performed on a Bruker AVANCEIII HD 600 MHz spectrometer equipped with a triple resonance cryoprobe. 128 scans were accumulated to acquire each spectrum.

*$^{19}F$ nuclear magnetic resonance (NMR) spectroscopy.* Protein samples used for $^{19}$F NMR analysis were diluted to concentrations of 75–280 μM in 500 μL of 10 mM potassium phosphate buffer (pH 6.0), 100 μM EDTA, 0.02% sodium azide, and 10% $D_2O$. All $^{19}$F NMR spectra were acquired with a Bruker Avance 500 MHz spectrometer with 10 sec acquisition delays. 512 scans were accumulated for one-dimensional $^{19}$F chemical shift analysis at 5, 10, 15, 20, 25, 30, and 35 °C. Data were processed with an exponential window function (20 Hz line-broadening) using TopSpin 3.6.1 (Bruker Biospin). The $^{19}$F NMR spectra of fluorinated HEX, VFIT, and VFIY showed two resonances, which we

interpret to correspond to two distinct conformations that do not rapidly exchange with each other on the $^{19}$F-NMR timescale. All spectra were deconvoluted into two Lorentzian curves using Python v3.7.9 with the LMFIT package for non-linear least-squares minimization and curve-fitting (*51*). The resulting two curves were integrated to measure the relative populations of each conformational state. The ratio of the two populations were used to calculate equilibrium constants at each temperature (Supplementary Table 8) and fit using the Solver function of Excel to the nonlinear van't Hoff equation (*52*) (Equation 1), where enthalpy and entropy are not assumed to be temperature-independent (i.e. $\Delta C_p \neq 0$) to extract thermodynamic parameters of equilibrium. When fitting to the linear van't Hoff equation, $\Delta C_P$ was simply set to zero.

$$\ln(K_{eq}) = -\frac{\Delta H^{\circ}_{ref}}{R}\left(\frac{1}{T}\right) + \frac{\Delta S^{\circ}_{ref}}{R} - \frac{\Delta C_P}{R}\left[\left(\frac{T - T_{ref}}{T}\right) + \ln\left(\frac{T_{ref}}{T}\right)\right]$$

**Equation 1. Nonlinear van't Hoff equation**. $T_{ref}$ is an arbitrary reference temperature (set to 298 K), $\Delta H^{\circ}_{ref}$ and $\Delta S^{\circ}_{ref}$ are $\Delta H^{\circ}(T)$ and $\Delta S^{\circ}(T)$ evaluated at $T_{ref}$, respectively.

*Crystallization.* Purified AAT variants were prepared in 20 mM potassium phosphate buffer (pH 7.5) with 2 mM EDTA and 10 uM PLP to a final concentration indicated on Supplementary Table 5. Maleic acid was dissolved in water to make a 1 M stock solution, and then added to each protein solution yielding a final maleate concentration of 20 mM. For each enzyme variant, we carried out initial crystallization trials in 15-well hanging drop format using EasyXtal crystallization plates (NeXtal) and a crystallization screen that was designed to explore parameter space around the crystallization conditions reported by Islam *et al.* (*15*). Crystallization drops were prepared by mixing 1 μL of protein solution with 1 μL of the mother liquor and sealing the drop inside a reservoir containing 500 μL of mother liquor. The mother liquor solutions contained ammonium sulfate as a precipitant and the specific growth conditions that yielded the crystals used for X-ray data collection are provided in Supplementary Table 5. For all six enzymes, a microseeding protocol was required to obtain high-quality crystals. Microseeds were prepared by crushing initial crystals in their mother liquor using a glass rod, and were subsequently streaked into the crystallization drops using a cat whisker. Because maleate was required for crystallization, ligand-free structures were obtained by soaking the crystal in new drops of 1 μL mother liquor containing 10 μM PLP but no maleate, allowing maleate in the crystal to diffuse out. Each crystal was treated in this way six subsequent times, 12 hours apart, to achieve removal of bound maleate (Supplementary Figure 9).

*X-ray data collection and processing.* Prior to X-ray data collection, crystals were mounted on polymer MicroMounts (MiTeGen) and sealed using a MicroRT tubing kit (MiTeGen). Single-crystal X-ray diffraction data was collected on beamline 8.3.1 at the Advanced Light Source. The beamline was equipped with a Pilatus3 S 6M detector and was operated at a photon energy of 11111 eV. Crystals were maintained at either 100 K, 278 K, or 303 K throughout the course of data collection. Each data set was collected using a total X-ray dose of 50–100 kGy and covered a 180° wedge of reciprocal space. Multiple data sets were collected for each enzyme variant either from different crystals, or if their size permitted, from unique regions of single large crystals.

X-ray data were processed with the Xia2 program (*53*), which performed indexing and integration with DIALS (*54*), followed by scaling with DIALS.SCALE (*55*). The resolution cut-off was taken where the $CC_{1/2}$ and $\langle I/\sigma I \rangle$ values for the intensities fell to approximately 0.5 and 1.0 respectively.

*Structure determination.* We obtained initial phase information for calculation of electron density maps by molecular replacement using the program Phaser (*56*), as implemented in v1.17.1.3660 of the

PHENIX suite (*57*), with the crystal structure of wild-type *Escherichia coli* AAT complexed with *N*-phosphopyridoxyl-L-glutamic acid (PDB ID: 1X28 (*15*)) as search model. All AAT variants crystallized in the same crystal form, containing two chains of the molecule in the crystallographic asymmetric unit. Next, we rebuilt the initial model using the electron density maps calculated from molecular replacement. We then performed additional, iterative refinement of atomic positions, individual atomic displacement parameters (B-factors), and occupancies using a translation-libration-screw (TLS) model, a riding hydrogen model, and automatic weight optimization, until the model reached convergence. All model building was performed using Coot 0.8.9.2 (*58*) and refinement steps were performed with phenix.refine (v1.17.1.3660) within the PHENIX suite (*57, 58*). Restraints for PLP in its internal aldimine form were generated using phenix.elbow (*59*), starting from coordinates available in the Protein Data Bank (*60*) (PDB ligand ID: PLP), and manually edited to ensure planarity of the pyridine ring and correct geometry of the imine bond between PLP and K246 (restraints available as Supplementary Data Files). Further information regarding model building and refinement, as well as PDB accession codes for the final models, are presented in Supplementary Table 6 and Supplementary Table 9.

**Acknowledgements**

R.A.C. acknowledges an Early Researcher Award from the Ontario Ministry of Economic Development & Innovation (ER14-10-139), and grants from the Natural Sciences and Engineering Research Council of Canada (RGPIN-2016-04831) and the Canada Foundation for Innovation (26503). The authors thank Michael D. Toney for providing the wild-type *E. coli* AAT expression vector. Beamline 8.3.1 at the Advanced Light Source is operated by the University of California San Francisco with generous support from the National Institutes of Health (R01 GM124149 for technology development and P30 GM124169 for user support), and the Integrated Diffraction Analysis Technologies program of the US Department of Energy Office of Biological and Environmental Research. The Advanced Light Source (Berkeley, CA) is a national user facility operated by Lawrence Berkeley National Laboratory on behalf of the US Department of Energy under contract number DE-AC02-05CH11231, Office of Basic Energy Sciences.

### Author Contributions

A.D.S. and R.A.C. conceived the project. A.D.S. performed computational design experiments. A.D.S. and M.G.E. purified proteins. A.D.S., S.M.F., and S.T.K. performed mutagenesis and cloning experiments. A.D.S. performed enzyme kinetics experiments. A.D.S. and A.M.D. performed NMR experiments. A.D.S. and N.K.G. designed NMR experiments. J.M.R. and M.C.T. crystallized proteins and performed X-ray diffraction experiments. R.A.C. and J.M.R. performed refinements. M.C.T. designed X-ray crystallography experiments. R.A.C. and A.D.S. wrote the manuscript. N.K.G., A.M.D., and M.C.T. edited the manuscript.

### Competing Interests

The authors declare no competing interests.
